# Transcriptomic analysis of genotypes derived from *Rosa wichurana* unveils molecular mechanisms associated with quantitative resistance to *Diplocarpon rosae*

**DOI:** 10.1101/2025.06.13.659350

**Authors:** L. Lambelin, T. Thouroude, J. Jeauffre, J. Chameau, C. Vilfroy, C. Boursier, S. Aubourg, S. Pelletier, D. C. Lopez Arias, L. Hibrand-Saint Oyant, V. Soufflet-Freslon, F. Foucher, S. Paillard

**Affiliations:** IRHS-UMR1345, Université d’Angers, INRAE, Institut Agro, SFR 4207 QuaSaV, 49071 Beaucouzé, France; ANAN, SFR 4207 QuaSaV, 49071 Beaucouzé, France; University of Minnesota, Saint Paul, MN, USA

**Keywords:** *Rosa*, black spot disease, defense response, QTL, RNA-seq

## Abstract

Black spot disease, caused by the hemibiotrophic fungus *Diplocarpon rosae*, is a major foliar disease of garden roses. Resistant cultivars offer an alternative to fungicides, but the genetic basis of resistance is not well known. Understanding these mechanisms is essential for the effective and durable deployment of resistant cultivars. A hybrid of *Rosa wichurana* (RW) exhibits quantitative resistance to black spot. Analysis of an F1 progeny from a cross between RW and the susceptible cultivar *Rosa chinensis* ‘Old Blush’ showed that RW’s resistance mainly involves two Quantitative Trait Loci (QTLs), on linkage groups 3 (B3) and 5 (B5). This study aimed to better characterize the molecular mechanisms underlying these QTLs. RNA sequencing was performed on inoculated and non-inoculated samples of RW and four F1 genotypes with different QTL combinations, at 0, 3, and 5 days post-inoculation (dpi). The comparison of inoculated and non-inoculated samples revealed that most genes were differentially expressed at 3 dpi. For all the genotypes, we observed shifts in expression of genes involved in plant defense, as well as genes participating in hormonal pathways and calcium-mediated signaling. Genotypes harboring QTL B3 showed “classic” defense responses involving pathogen recognition, signaling, ROS production, callose deposition, and localized cell death. In contrast, QTL B5 was associated with few shared DEGs, making it harder to define its role. The quantitative nature of RW’s resistance may result in this complex regulation of gene expression.

**Key message:** Two QTLs derived from *Rosa wichurana* contribute differently to rose black spot resistance, with QTL B3 linked to classical defense responses and QTL B5 suggesting a more complex regulatory mechanism.

## Introduction

Black spot disease is a major fungal disease of garden roses worldwide. Caused by the hemibiotrophic ascomycete *Diplocarpon rosae* (anamorph of *Marssonina rosae*), obligate to *Rosa* genus, this foliar disease manifests from spring to autumn. The symptoms include dark, rounded spots with fringed margins on the upper side of leaves, chlorosis and defoliation, which considerably decrease the aesthetic value of rose bushes. In highly susceptible cultivars, the resulting reduced vigor can lead to increased vulnerability to other stresses and potential plant death (Gachomo and Kotchoni 2007). While black spot can be managed with fungicides, their use is decreasing for several reasons. First, the growing awareness of their risks for health and environment led to their restrictions in Europe (Zlesak 2007; Leus 2017). In France, this directive is enforced through the Ecophyto Plans (Ecophyto, Ecophyto II, Ecophyto II+ and Ecophyto 2030), resulting in a ban on pesticide use in public gardens since 2017 and private gardens since 2019 (Labbé 2014). Moreover, from an evolutionary point of view, chemical management of pests and diseases favors the selection of pesticide-resistant organisms, making this approach non-durable. Last, consumers tend to prefer low-maintenance roses that can grow almost independently and do not require chemical treatments (Waliczek et al. 2015). Consequently, the demand for disease-resistant roses has surged.

Despite progress over the past 30 years in breeding new varieties with enhanced black spot resistance, significant challenges remain. Indeed, roses are perennial bushes and are propagated vegetatively. Therefore, understanding the underlying processes of resistance is crucial to identify genotypes with durable resistances, i.e., resistances that will not be easily overcome by pathogens. To date, six major resistance genes (*Rdr1* to *Rdr6*) have been discovered in roses (von Malek and Debener 1998; Hattendorf et al. 2004; Whitaker et al. 2010a; Zurn et al. 2018; Lopez Arias et al. 2023; Moore et al. 2023). *Rdr1* and *Rdr2* originate from *Rosa multiflora* and are located on chromosome 1 (von Malek and Debener 1998; Hattendorf et al. 2004), *Rdr3* was inherited from chromosome 6 of ‘George Vancouver’ (Zurn et al. 2020), *Rdr4* from chromosome 5 of Brite Eyes^TM^ (Zurn et al. 2018), *Rdr5* from chromosome 7 of Ramblin’ Red® (Lopez Arias et al. 2023), and *Rdr6* from chromosome 5 of Baby Love™ (Moore et al. 2023). These genes are associated with race-specific resistances, however, their range of action varies: *Rdr2* confers immunity against race 4 (Hattendorf et al. 2004; Whitaker et al. 2010b), while all five others confer broad-spectrum resistance against several *D. rosae* isolates and races (von Malek and Debener 1998; Hattendorf et al. 2004; Whitaker et al. 2010a; Zurn et al. 2018; Lopez Arias et al. 2023; Moore et al. 2023). Currently, 14 pathogenic races of *D. rosae* have been documented using a set of 10 differential hosts (Whitaker et al. 2010b; Rouet et al. 2020; Zlesak et al. 2020). Recently, several Quantitative Trait Loci (QTLs) for black spot resistance have been identified in roses, exhibiting either major or minor effects and located across all seven rose chromosomes (Lopez Arias et al. 2020; Rawandoozi et al. 2022; Lau et al. 2023). Despite having a lower effect compared to major genes, quantitative resistance remains a valuable resource for breeding. Indeed, resistances relying on multiple QTLs are often considered more durable than those relying on only one or two major genes (Poland et al. 2009; St Clair 2010). In addition to these genetic analyses, transcriptomic studies have been conducted to unravel the molecular mechanisms of the rose*-D. rosae* interaction (Neu et al. 2019; Song et al. 2024). Both studies unveiled common sets of differentially expressed genes (DEGs), such as pathogenesis-related (PR) proteins, defense-related transcription factors (WRKY), and genes involved in flavonoid and phenylpropanoid biosynthesis. They also identified DEGs that could be genotype-specific, particularly in hormone signaling pathways. However, these studies were not associated with a genetic analysis, and the genetic basis of the cultivars used is not known.

The present work aimed to investigate the quantitative resistance to black spot disease in a hybrid genotype derived from *Rosa wichurana*, a species that has been frequently used in breeding programs (Byrne et al. 2007). A previous genetic study based on an F1 progeny (OW) resulting from the cross between the susceptible *Rosa chinensis* ‘Old Blush’ (OB) and this *R. wichurana* hybrid (RW) identified two major resistance QTLs on chromosomes B3 and B5, accounting for 22.1 % and 11% of the explained variance, respectively (Lopez Arias et al. 2020). These two QTLs were also found in two other F1 progenies derived from RW, and were consistently detected across years and locations. Our objective was to characterize the global defense response associated with each of these QTLs, at the molecular level. In this framework, F1 genotypes with different QTL combinations (either QTL B3, QTL B5, or both B3 and B5) have been chosen. Using RNA sequencing, their transcriptome was captured during the first stages of infection, 0, 3, and 5 days post-inoculation (dpi). The comparison of inoculated and non-inoculated (mock) samples at each time point yielded DEGs during the host-pathogen interaction. Common DEGs between the individuals harboring a common QTL were compared and revealed candidate mechanisms for black spot resistance associated with QTLs B3 and B5.

## Material and methods

### Plant material

Based on the genotypic data of individuals from the OW progeny, four individuals with different QTL combinations were selected (**Fig. 1** and **Table 1**). The objective was to focus on the two major QTLs found in RW’s genome (QTL B3 and QTL B5). Therefore, individuals harboring other minor QTLs were excluded (i.e., QTL B4 inherited from RW and QTL A1 inherited from OB, as described by Lopez Arias et al. 2020). Moreover, the confidence intervals (CIs) were highly variable from one year to another for QTLs B3 and B5, and the authors suspected the existence of two QTLs on chromosome B5 (Lopez Arias et al. 2020). Therefore, we decided to consider a top (B5A) and a bottom (B5B) CI for QTL B5. The QTL CIs were defined as follows:

- QTL B3: from 4 to 28.3 cM, it included all annual QTL peaks and the CI calculated from the average over the years;
- QTL B5A: from 6 to 22.1 cM, it included the CIs of the QTLs detected in 2017 and 2018, which overlapped perfectly and had the same peak, and the CI calculated from the average over the years;
- QTL B5B: from 22.1 to 48.3 cM, it included the end of the CI of the QTL detected in 2016, the year when the LOD curve clearly revealed two QTLs on chromosome B5.

**Fig. 1.**
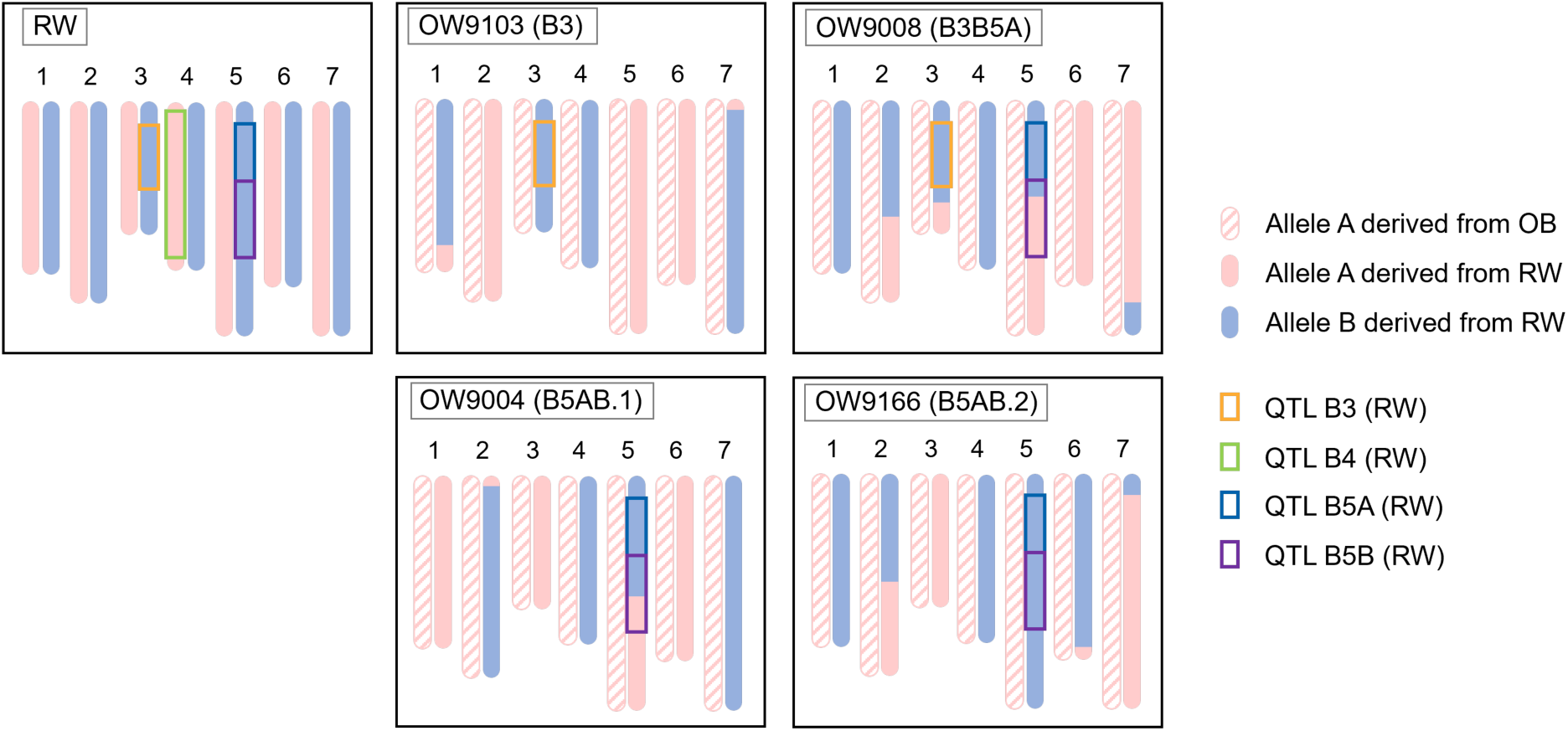
Schematization of the genotyping of RW and the four OW individuals. Individual names are followed by their genotypic class in parentheses. Chromosomes are represented by vertical bars. Maternal and paternal alleles are represented in color, QTLs are represented by colored boxes according to the results from Lopez Arias et al. (2020) using OW progeny

**Table 1.**
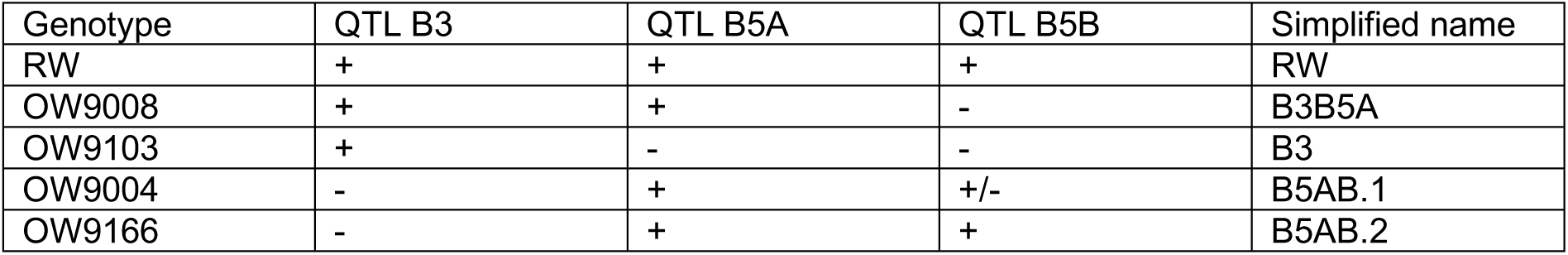
Summary of the five rose genotypes used in the RNA-seq assay. ‘+’ and ‘-’ refer respectively to the presence or absence of the QTL.

For ease of comprehension, the individuals will be referred to as their QTL combination, i.e., B3B5A (OW9008), B3 (OW9103), B5AB.1 (OW9004) and B5AB.2 (OW9166) (**Fig. 1** and **Table 1**). The resistant male parent RW was also included in the RNA-seq for comparison.

### Greenhouse assay

The assay was performed in May 2023. Vegetatively propagated clones of each genotype were grown in pots of diameter 10.5 cm in a greenhouse at 21°C with a 16-hour photoperiod. Four-month-old plants were divided into three blocks following a double split-plot design. Each block was spatially separated into two sub-blocks between the inoculated condition (I) and the non-inoculated condition (N). Each sub-block comprised three rows corresponding to the dpi when plants were sampled. Plots of three clones per genotype were randomized within rows, and the order of the rows within a sub-block was randomized as well.

In each sub-block, a fourth row was not sampled and kept for later phenotyping. This design was chosen to avoid mistakes while sampling the rose clones.

For inoculum preparation, *D. rosae* strain DiFRA_67 (Marolleau et al. 2020) was propagated on cellophane sheets overlaid onto malt agar (10 g/L christomalt, 15 g/L agar) and incubated at 20°C for 10 to 15 days. Cellophane sheets were then conserved at -20°C before being used for the inoculation assay. Conidia were resuspended in water and the concentration was adjusted to 82,000 conidia/mL right before the experiment.

Inoculation took place according to the method described by Soufflet-Freslon et al. (2019). The inoculated group (I) was sprayed with a solution of conidia, and the non-inoculated control (N) was sprayed with distilled water. Groups of three vegetatively-propagated clones were sampled 30 min after inoculation (referred to as 0 dpi), then 3 and 5 dpi. A total of nine leaves (three from each clone) were pooled for each sample. Each block was sprayed at 24-hour intervals (first with water for the non-inoculated sub-block, then with the inoculum for the inoculated one) and used as biological replicates (referred to as repetitions A, B and C). Therefore, as each genotype was sampled under two conditions at three time points and in three replicates, a total of 90 samples were prepared for RNA-seq assay. Samples were coded as follows: Genotype_Treatment_dpiReplicate.

### Phenotypic scoring of symptoms

Three inoculated plants and three non-inoculated plants per genotype were used to assess the disease levels at 28 dpi. Symptoms were visually scored using a 0 (no symptom) to 10 (total defoliation) scale (**Table 2**). The same number of plants of the susceptible cultivar OB were used as controls for successful inoculation.

**Table 2.** Scoring scale for black spot disease of rose.

| Score | Description |
| --- | --- |
| 0 | No symptom |
| 1 | A few spots on 10 % of the foliage |
| 2 | > 10 % to 25 % of the foliage |
| 3 | > 25 % to 33 % of the foliage |
| 4 | > 33 % to 50 % of the foliage |
| 5 | > 50 % to 66 % of the foliage |
| 6 | > 66 % to 75 % of the foliage |
| 7 | > 75 % of the foliage, no defoliation |
| 8 | > 75 % of the foliage, defoliation |
| 9 | 100 % of the foliage, strong defoliation |
| 10 | Complete defoliation |

### RNA-seq assay

Whole leaves were collected and petioles were removed right before the samples were flash-frozen in liquid nitrogen, then stored at -80°C. Before RNA extraction, leaves were ground using a HG-400 MiniG^®^ grinder (SPEX^®^ SamplePrep). Total RNA was extracted with a NucleoMag™ RNA kit (Macherey-Nagel™, Düren, Germany) and IDEAL™ 32 extraction robot (Innovative Diagnostics, Grabels, France). A few adjustments were made to the manufacturer’s instructions: on step 1 of the protocol, MR1 buffer was replaced by a home-made lysis solution, composed of 400 µL LBP buffer, 8 mg PVP 10 and 7 µL TCEP for each sample; on step 3, 24 μL (instead of 28 µL) re-suspended NucleoMag™ B-Beads were added.

RNA concentration, possible degradation and contamination of the samples by other molecules were controlled with a Nanodrop™ One machine (Thermo Fisher Scientific, Waltham, MA, USA) and an Agilent Bioanalyzer (Agilent Technologies, Santa-Clara, CA, USA) system before sending the samples to the sequencing company.

All the samples were then sent to BGI Genomics (Varsaw, Poland), and the next steps of stranded library preparation and RNA sequencing were taken care by the company. Paired-end, 150 bp reads were obtained using DNBseq™ Sequencing Technology, with a mean sequencing depth of 50 M reads per sample for RW, and 25 M reads per sample for the rest of the samples.

### Genome sequence of RW and reference transcriptome

As a high-quality haplotype-resolved annotated genome sequence of RW was available (bioproject PRJEB88836 on the European Nucleotide Archive website), this sequence was used to generate a reference transcriptome to map the RNA-seq data. A customized haploid reference transcriptome was built using a reciprocal BLASTn procedure between the two haplotypes, with the *e-value* threshold set to 10^-20^. The first haplotype was then combined with the hemizygous genes found in the other haplotype. The reference transcriptome index was built using the corresponding mRNA sequences with Salmon v1.10.2 with default parameters (Patro et al. 2017).

### Quality controls and data analysis

#### Read trimming and mapping

Raw reads were trimmed using fastp (Chen et al. 2018), with the parameters -*D --dup_calc_accuracy 3 -g -x -z 4 -p -w 16 -q 30 -l 50 -i*. Their quality was assessed with MultiQC (Ewels et al. 2016). Clean reads were pseudo-mapped against the customized haploid reference transcriptome of RW using Salmon v1.10.2 with default parameters (Patro et al. 2017).

#### Fungal read count

Before analyzing the RNA-seq data, we wanted to make sure that mock samples were free from *D. rosae* contamination. In this framework, we compared the number of reads with high sequence similarity to the *D. rosae* genome between inoculated and non-inoculated samples as follows. The cleaned and paired forward sequences of each sample were aligned to the *D. rosae* draft genomic sequence (Neu et al. 2017) using BLASTn with default parameters and an *e-value* threshold of 10^-3^. For each read aligning with the *D. rosae* genome, the best BLAST hit was selected, and only reads with more than 90% identity over a minimum length of 140 nucleotides were considered. Reads aligned with *D. rosae* sequences coding for ribosomal RNA were eliminated.

#### Differential expression analysis

For each genotype, DEGs between inoculated and mock samples were compared for each time point of the interaction. A gene was considered as differentially expressed if its adjusted *p-value* (Benjamini-Hochberg method, 1995) was inferior than 0.05 and the absolute value of its log2 fold change (Lfc) above 0.58 (equivalent to a change of 1.5-fold in gene expression). Differential expression analysis was performed using DESeq2 package (Love et al. 2014) in R software version 4.4.0 (R Core Team 2024). As the three biological repetitions were sampled with 24-hour intervals, variability between repetitions was expected. To account for this effect, the design formula used for DESeq2 was: ‘design = ∼ repetition + condition’, with ‘repetition’ being the effect of the biological repetition (A, B or C), and ‘condition’ being either inoculated or mock.

#### Sample clustering

A Principal Component Analysis (PCA) was conducted using the DESeq2 package (Love et al. 2014) and the ggplot2 package (Wickham 2016) in R software version 4.4.0 (R Core Team 2024) to visualize sample distribution between genotypes. For each genotype, RNA-seq data were filtered based on the DEGs for at least one condition to calculate Pearson’s correlations between genotype samples. These analyses identified outliers that were removed. After eliminating these outliers, the differential expression analysis was re-run to ensure accurate results.

#### Analysis of gene functions and connection with the QTL class

To analyze the results of the differential analysis, we used the functional annotation of RW’s genome available at this time (bioproject PRJEB88836 on the European Nucleotide Archive website). This annotation used BLAST comparisons between genes of RW and the databases SwissProt and TAIR10. It came with a UniProt code or reference that allowed us to easily search for details on the functions of these genes in model plant species on UniProt database (The UniProt Consortium 2023).

Gene ontology (GO) enrichment analysis was done with AgriGO v2.0 (Du et al. 2010; Tian et al. 2017). For each genotype, a Single Enrichment Analysis (SEA) was performed for up- and downregulated genes, using the customized haploid transcriptome of RW. The parameters were the following: Fisher’s exact test, multi-test adjustment method of Benjamini-Yekutieli (2001) (False-discovery rate (FDR) under dependency), significance level of 0.05 and minimum number of mapping entries of 3.

The analysis of common genes for individuals sharing a QTL was done using the ‘intersect’ function in R and by filtering the genes only shared between individuals carrying QTL B3 or QTL B5 with Excel. Figures were generated using the ggplot2 package (Wickham 2016) in R software.

## Results

### Phenotyping

The success of the inoculation was assessed using the susceptible cultivar OB, with all inoculated plants showing symptoms and having a high disease score (**Fig. 2**). As expected for a resistant genotype, inoculated plants of RW showed very few symptoms, and their scores ranged from 0 to 1. As for the inoculated F1 progenies, they exhibited intermediary phenotypes, with moderate levels of resistance.

**Fig. 2.**
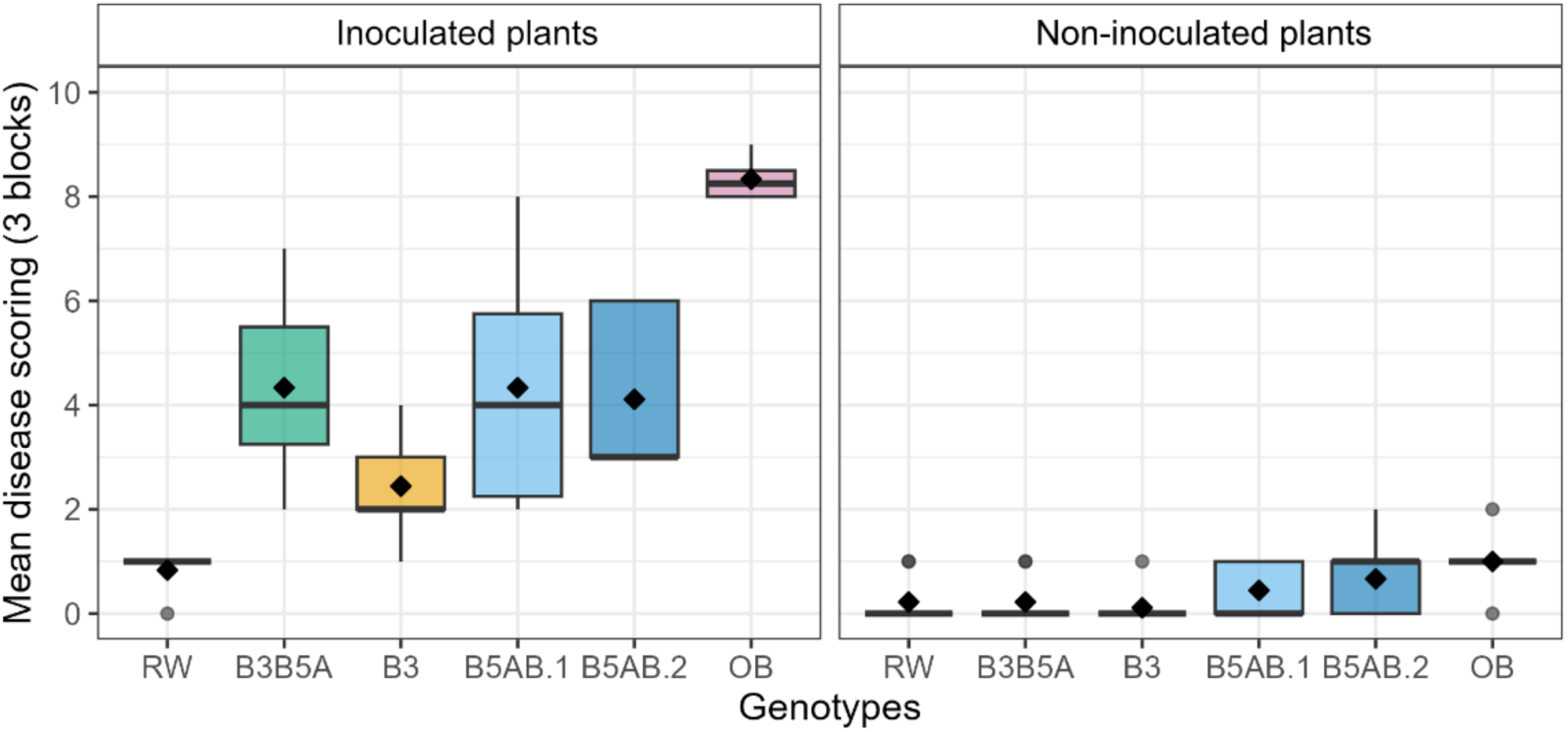
Mean disease scoring of the six genotypes at 28 dpi, for the three biological repetitions (blocks), for inoculated plants and non-inoculated plants. The bold line represents the median and the diamond represents the mean

Slight symptoms were observed on the mock plants (**Fig. 2)**, especially for B5AB.1, B5AB.2 and OB. Therefore, to verify that the RNA-seq control samples were not contaminated with *D. rosae*, a count of fungal reads was performed for both inoculated and mock samples (**Online Resource 1**). On average, fungal reads per inoculated samples represented around 0.38 % of the total reads. Fungal reads per mock sample did not excess 0.01 %, except for samples RW_N_0C (0.662 %), OW9103_N_5A (0.011 %), OW9103_N_5B (0.016 %) and OW9166_N_5_A (0.016 %). Concerning sample RW_N_0C, the significant fungal read number probably stemmed from an error during sample manipulation and did not originate from contamination in the greenhouse. Therefore, this sample was removed from the analysis. For the three other samples, as their fungal read percentages were still much lower than the ones of inoculated samples, we chose to retain them for RNA-seq analysis.

### Quality control and alignment of transcriptome data

Raw transcriptomic data are available on the European Nucleotide Archive website under bioproject PRJEB90172. Out of the 90 samples collected from the four F1 and RW, two were lost during the RNA extraction process (OW9004_N_0A and OW9103_I_0A). All the remaining samples were successfully sequenced and resulted in a total of 1.1 Tb of cleaned data, with an average of 124,794,687 reads per library after filtering for RW, and 62,770,975 reads per library after filtering for the rest of the samples. In this dataset, the mean quality score was high and stable along the read length and the average GC content was 47.7 % (**Online Resource 2**). Therefore, the sequenced reads were of high quality and met the criteria of subsequent analyses.

The customized reference transcriptome resulted in a total of 76,434 genes: 53,716 genes from Hap1 and 22,178 hemizygous genes from Hap2. The average alignment rate of clean reads on the reference transcriptome was 80.7 % (**Online Resource 2**).

Sample distribution between genotypes was investigated with a PCA, revealing that samples clustered into five groups according to the genotype (**Online Resource 3**). Two samples (RW_N_5C, and OW9166_N_0C) were identified as technical outliers and removed from the analysis. Indeed, they did not cluster with the other samples of their respective genotypes and exhibited different gene expression.

Further, for each genotype, correlations between samples were investigated after filtering by genes that were differentially expressed between mock and inoculated samples in at least one time point (**Online Resource 4**). In general, time points 0 and 5 dpi were not well separated and often exhibited close correlation coefficients (R). By contrast, samples at 3 dpi usually clustered together and were less well correlated with samples at 0 and 5 dpi. Overall, the correlation coefficients between samples at 0 and 5 dpi and between samples at 3 dpi were higher than 0.85. Three samples (OW9004_I_3A, OW9103_I_3A and OW9166_I_3A) were different from their other replicates (R < 0.82 with the two other replicates) and were removed from analysis (**Online Resource 4**).

### Contrasted responses between genotypes

DESeq2 analysis on the 82 remaining samples showed that, in general, very few genes were differentially expressed during the first 30 min of interaction (inoculated *vs* mock samples at 0 dpi), irrespective of the genotype. The same phenomenon was observed for 5 dpi (**Fig. 3**). In consequence, very few DEGs were shared between the three time points (data not shown) and the temporal comparison of DEGs will not be further explored.

**Fig. 3.**
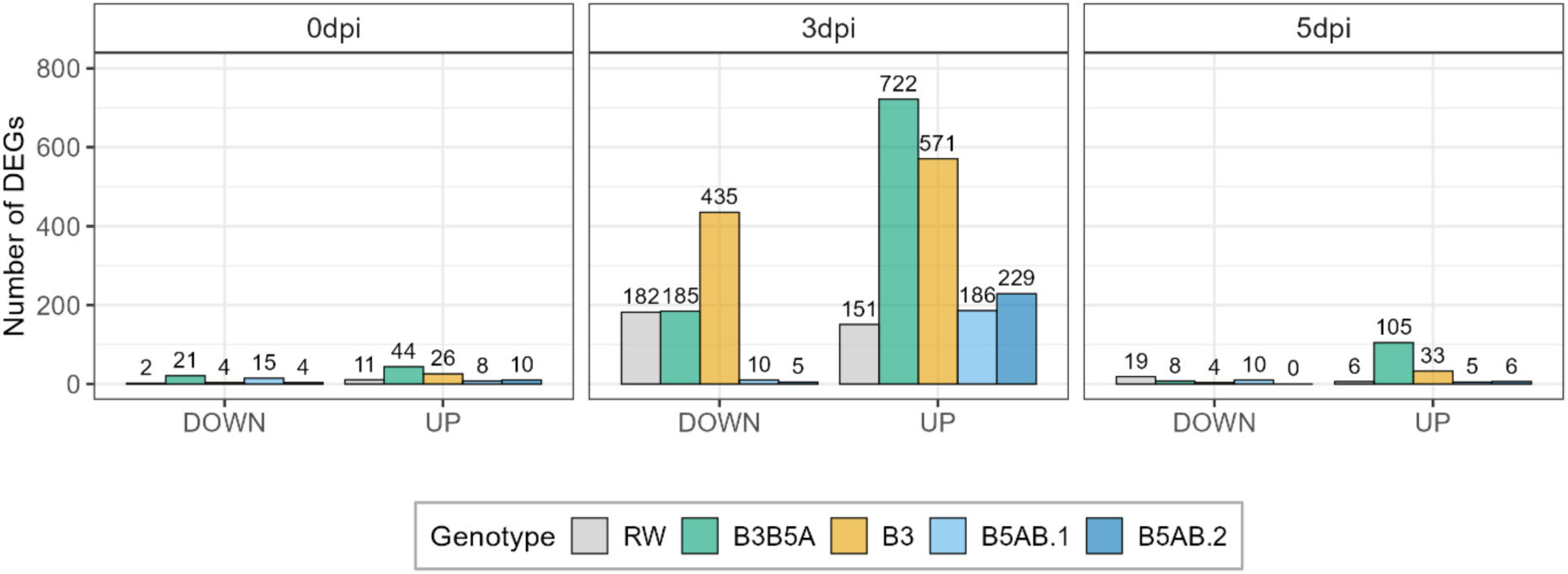
Number of DEGs for each time point (inoculated *vs* mock samples), for each genotype

By contrast, 3 dpi seemed to be the most interesting time point for this comparison study, as the most DEGs were found then: a total of 333 DEGs for RW, 907 for B3B5A, 1006 for B3, 196 for B5AB.1 and 234 for B5AB.2 (**Fig. 3**).

A GO enrichment analysis was performed for each genotype using AgriGO, then the enriched terms were compared. The objective of this study was to identify common processes between genotypes harboring the same major QTLs. Therefore, GO enrichment results were filtered based on the appearance of a term in at least two genotypes sharing a QTL (**Fig. 4**). The complete list of enriched GO terms is available in **Online Resource 5**. A GO term enriched in up- or downregulated genes in several genotypes does not always account for the same underlying genes, and some common DEGs do not belong to GO terms significantly enriched in up- or downregulated genes. Therefore, further comparisons of the common DEGs between genotypes are provided in the last part of the results.

**Fig. 4.**
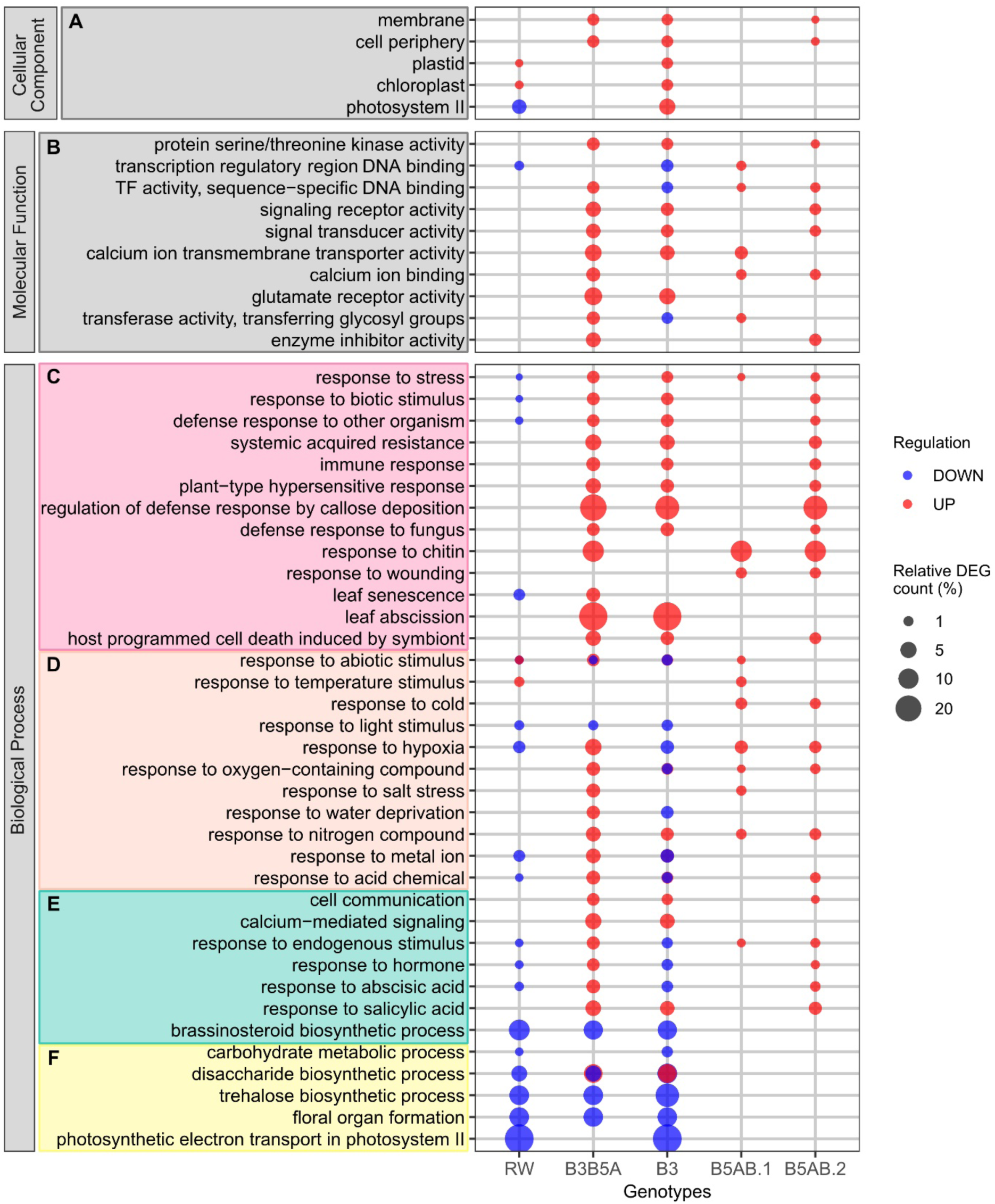
Gene Ontology (GO) enrichment bubble plots for each genotype at 3 dpi. Only the most relevant, significantly enriched terms are shown (FDR < 0.05); all terms are available in the **Online Resource 5**. Gene Ontology aspects: Cellular Component (A); Molecular Function (B) and Biological Process. Terms associated with biological processes are grouped and colored per family of GO terms: response to biotic stress (C), response to abiotic stress (D), signaling (E) and resource allocation and plant growth (F). Circle size indicates the number of DEGs per GO term relative to the number of genes per GO term in the reference annotation. Circle color indicates whether the term is associated with upregulation or downregulation

#### Singular profile of RW

RW exhibited a different GO enrichment pattern compared to the F1 genotypes. Indeed, RW exhibited few enriched GO terms, and terms that were associated with upregulated genes in the F1 genotypes were either downregulated in RW or were not significantly enriched (**Fig 4**). For example, RW showed a downregulation of the terms “response to stress”, “response to biotic stimulus”, “defense response to other organism” and “leaf senescence”, and the terms associated with a response to a fungus or chitin were not significantly enriched (**Fig. 4 C**). Moreover, terms related to signaling were downregulated or not enriched (**Fig. 4 B, E**). Globally, the defense responses in RW did not seem to be activated at 3 dpi.

#### Non-QTL-specific GO enrichment profiles

At first glance at **Fig. 4**, it appeared that the four F1 genotypes shared numerous GO terms, in particular B3B5A, B3 and B5AB.2, while B5AB.1 lacked some of these terms. The associated genes were generally upregulated, with exceptions in the response to abiotic stresses, hormones and transcription factor (TF) activity for B3, which were associated with gene downregulation (**Fig. 4 A, D, E**).

The cellular components “membrane” and “cell periphery” were enriched in upregulated genes in B3B5A, B3 and B5AB.2 (**Fig. 4 A**). Concerning molecular functions, the genes associated with the term “transcription regulatory region DNA binding” were downregulated in RW and B3, but upregulated in B5AB.1. As for “TF activity, sequence-specific DNA binding”, genes were upregulated in B3B5A, B5AB.1 and B5AB.2, but downregulated in B3 (**Fig. 4 B**). The “signaling receptor activity” and “signal transducer activity” genes were upregulated in B3B5A, B3 and B5AB.2 (**Fig. 4 B**). Genes related to calcium ion-mediated signaling were upregulated in the four F1, with “calcium ion binding” being upregulated in B3B5A, B5AB.1 and B5AB.2, and “calcium ion transmembrane transporter activity” upregulated in B3, B3B5A and B5AB.1 (**Fig. 4 B**). As for biological processes, the GO term “response to stress” was highly significantly enriched in upregulated genes in the four F1 genotypes, B3B5A, B3, B5AB.1 and B5AB.2 (**Fig. 4 C**). However, the genes associated with the GO terms “response to biotic stimulus”, “defense response to other organism”, “immune response”, “plant-type hypersensitive response”, “defense response to fungus”, “regulation of defense response by callose deposition” and “host programmed cell death induced by symbiont” were upregulated only in B3B5A, B3 and B5AB.2 (**Fig. 4 C**). Responses to various abiotic stimuli were differentially expressed and differed among the genotypes. The genes involved in “response to abiotic stimulus” were upregulated in B5AB.1, and both upregulated and downregulated in RW, B3B5A and B3, but was not significantly enriched in B5AB.2, despite other related terms being enriched. The “response to hypoxia” genes were downregulated in RW and B3 but upregulated in B3B5A, B5AB.1, and B5AB.2 (**Fig. 4 D**). The genes linked to “response to nitrogen compound” GO term were significantly upregulated in the four F1 genotypes. The genes involved in “response to hormone” and/or “hormone-mediated signaling pathway” were downregulated in RW and B3, but “response to hormone” genes were upregulated in B3B5A and B5AB.2 (**Fig. 4 E**). Different hormonal pathways were significantly enriched in RW, B3B5A, B3 and B5AB.2, but none in B5AB.1. Notably, “response to abscisic acid” genes were downregulated in RW and B3, but upregulated in B3B5 and B5AB.2. In contrast, “response to salicylic acid” genes were upregulated in B3B5, B3 and B5AB.2, but not significantly enriched in RW (**Fig. 4 E**).

#### QTL-specific GO terms

In the F1 genotypes harboring QTL B3 (i.e., B3B5A and B3), the infection by *D. rosae* triggered transcriptional modifications in photosynthetic pathways, brassinosteroid biosynthetic pathways, sugar metabolism and organ growth. Genes with functions related to chloroplasts and plastids were upregulated in RW and B3. Terms referring to photosystems were also enriched, but these genes were downregulated in RW and upregulated in B3 (**Fig. 4 C**). The genes involved in “photosynthetic electron transport in photosystem II” were downregulated in both RW and B3 (**Fig. 4 F**), while those related to “photosystem II” was downregulated in RW but upregulated in B3 (**Fig. 4 C**). For B3B5A, “photosystem II stabilization” and “photosystem II assembly” genes were downregulated, whereas “photosynthetic acclimation” genes were upregulated in B5AB.1 (**Online Resource 5**). In addition, the “response to light stimulus” genes were downregulated in RW, B3, and B3B5A (**Fig. 4 D**). Regarding the response to stress, genes involved in “leaf abscission” were only upregulated in B3 and B3B5A. In particular, the “brassinosteroid biosynthetic process” genes were downregulated in RW, B3, and B3B5A (**Fig. 4 E**), and was not found in B5AB.1 nor B5AB.2. Genes associated with carbohydrates and saccharides (e.g. “disaccharide biosynthetic process”) were downregulated in RW, and both downregulated and upregulated in B3B5A and B3 (**Fig. 4 F**). Finally, genes involved in floral organ formation were downregulated in RW, B3B5A and B3 (**Fig. 4 F and Online Resource 5**).

As for F1 genotypes harboring QTLs on chromosome B5 (i.e., B3B5A, B5AB.1 and B5AB.2), they did not share enriched GO terms exclusively with RW. Except for the term “response to cold”, no terms were only shared between B5AB.1 and B5AB.2. Therefore, it was not possible to identify QTL B5B-specific GO terms. Regarding stress response, B3B5A, B5AB.1 and B5AB.2 reacted to *D. rosae* with more precise GO terms such as “response to chitin”, and “response to wounding” was found for B5AB.1 and B5AB.2 (**Fig. 4 C**). Genes involved in salt stress response were only upregulated in B3B5A and B5AB.1.

### DEGs and candidate mechanisms for each QTL at 3 dpi

To get more insight into the molecular mechanisms underlying each QTL, the common DEGs at 3 dpi between genotypes sharing a QTL were investigated.

The objective of this section was to find DEGs shared between genotypes having a common QTL. As RW exhibited a singular expression pattern and shared few or no DEGs with the F1 genotypes having a common QTL (**Fig. 5**), the common DEGs for a QTL class were defined as follow: for QTL B3, genes that were differentially expressed in individuals B3 and B3B5A but not in B5AB.1 nor B5AB.2; for QTL B5A, genes that were differentially expressed in individuals B3B5A, B5AB.1 and B5AB.2 but not in B3; and for QTL B5B, genes that were differentially expressed in individuals B5AB.1 and B5AB.2 but not in B3 nor B3B5A. Assumptions on gene functions were based on RW genome annotation and Uniprot database (The UniProt Consortium 2023).

**Fig. 5.**
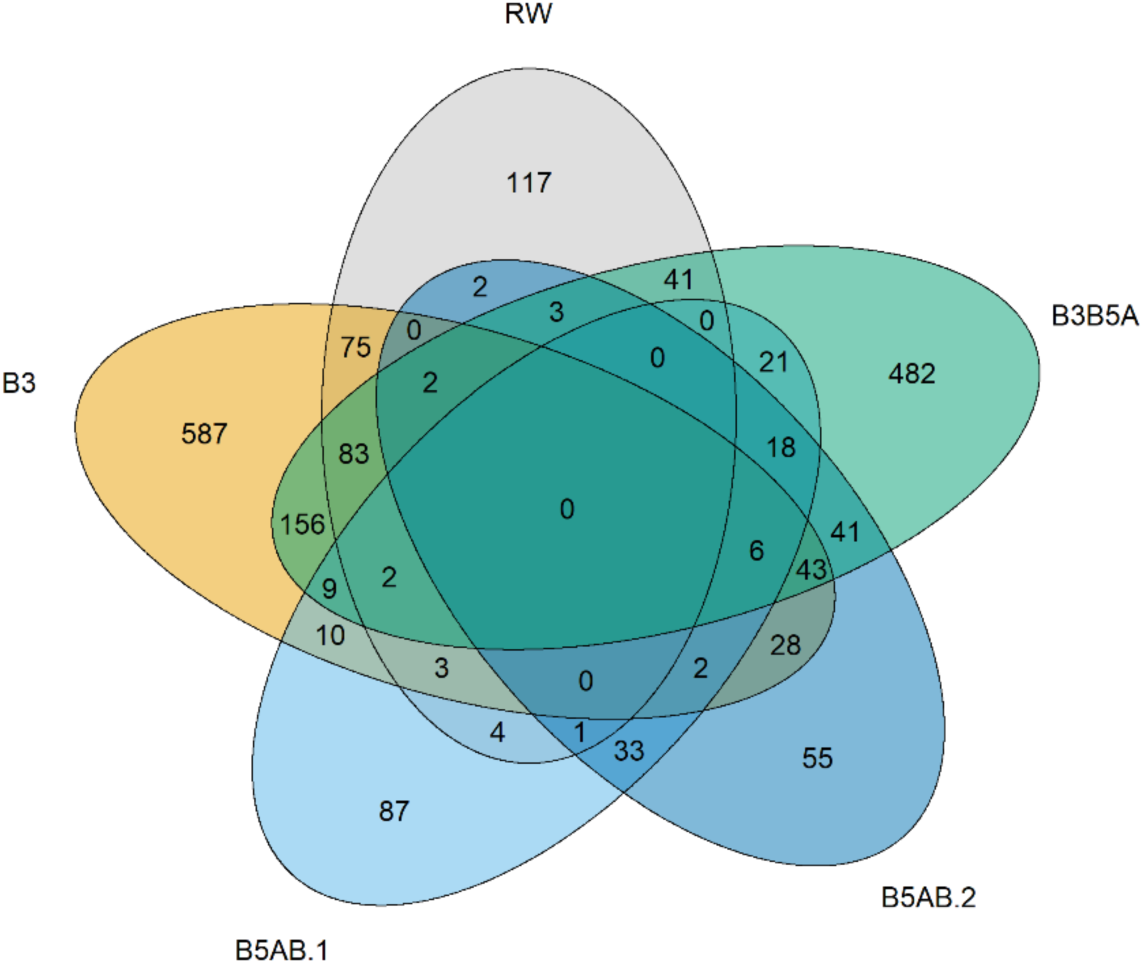
Shared DEGs (both upregulated and downregulated) at 3 dpi between the five genotypes

#### DEGs associated with QTL B3

Among the 301 common DEGs between B3B5A and B3, 62 were also shared with B5AB.1, B5AB.2 or both (**Fig. 5**). These genes shared between the four F1 genotypes may not be specific to one QTL or another. Let aside those genes, 239 DEGs were shared only between B3B5A and B3 (**Fig. 5**), of which 73 were downregulated, 155 were upregulated and 11 exhibited contrary expression patterns. Twelve genes lacked either a functional annotation or GO terms and their function could not be inferred (**Online Resource 6**). **Fig. 6** presents some of the common DEGs between B3B5A and B3 that could play a role in defense. Their differential expression in RW is also displayed although most genes were not differentially expressed (white cells).

**Fig. 6.**
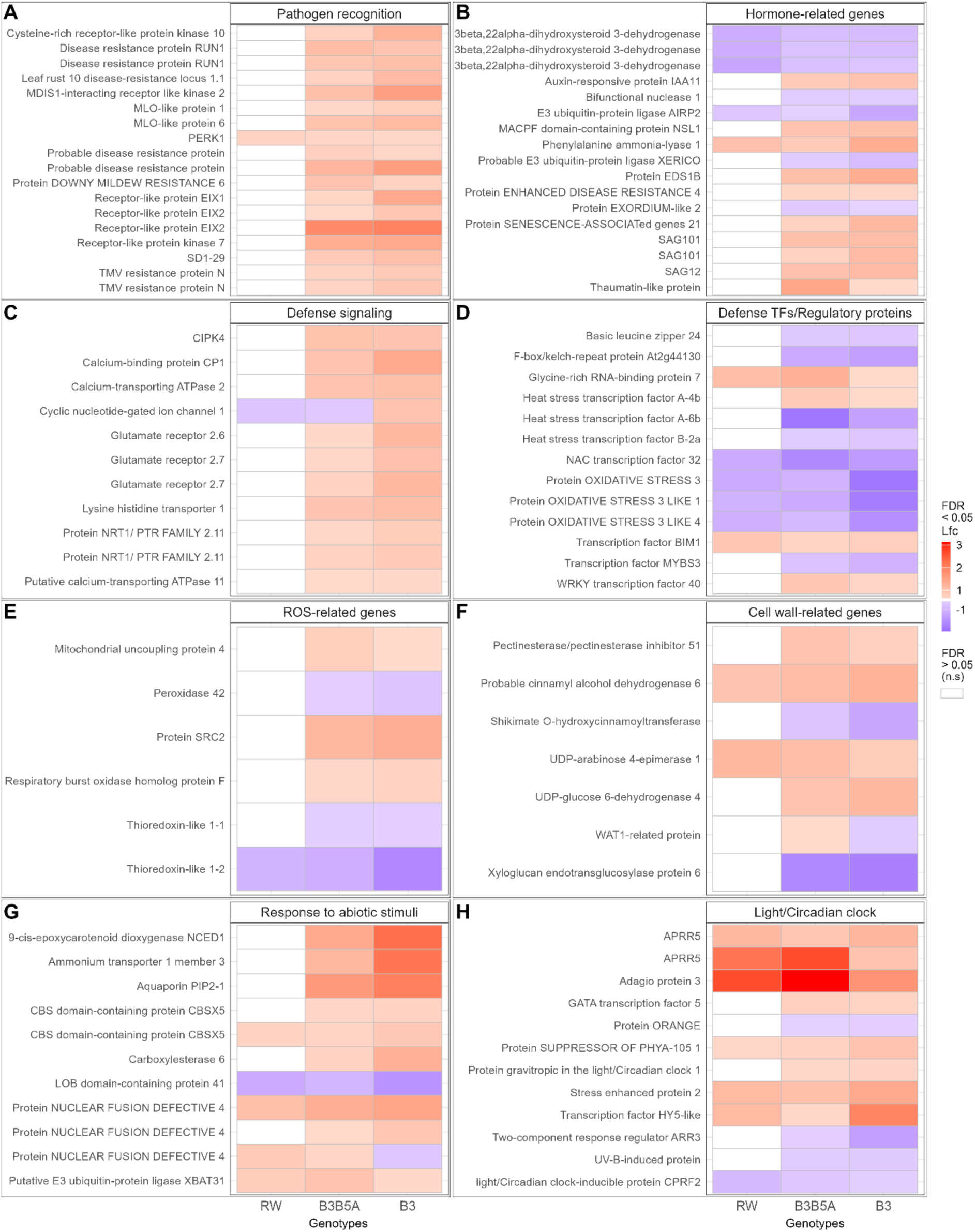
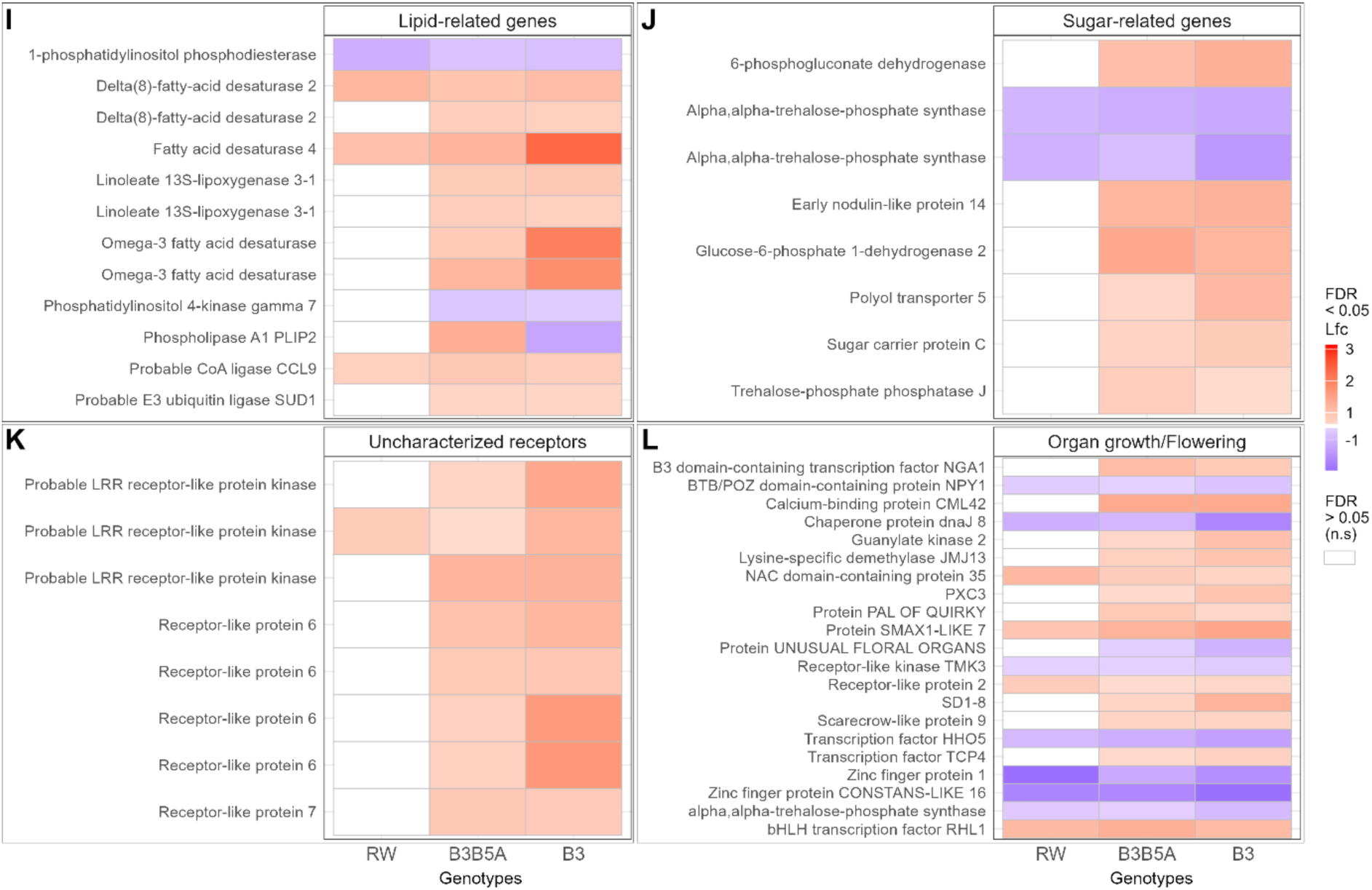
Heatmaps of annotated common DEGs between RW, B3B5A and B3 at 3 dpi. DEGs were classified according to their associated keywords or gene families. Genes not significantly (n.s.) differentially expressed are displayed in white. Fold changes, compared to the non-inoculated leaves, of down- (blue) and upregulation (red) are shown on a log2 scale (lfc). (A) Pathogen recognition; (B) Hormone-related genes; (C) Defense signaling; (D) Defense TFs/Regulatory proteins; (E) ROS-related genes; (F) Cell wall-related genes; (G) Response to abiotic stimuli; (H) Light/Circadian clock; (I) Lipid-related genes; (J) Sugar-related genes; (K) Uncharacterized receptors; (L) Organ growth/ Flowering

Many upregulated genes shared between B3B5A and B3 were related to defense response. These included eight genes annotated as disease-resistance proteins, five genes annotated as RLKs and three annotated as RLPs involved in response to pathogens, as well as two mildew resistance locus-O (MLO)-like proteins, a thaumatin-like protein, a probable WRKY 40 TF and a glycine-rich RNA-binding protein (**Fig. 6 A, D**). All these genes possessed GO terms related to a response to fungi or with a general defense process against pests and pathogens. Moreover, eight additional genes encoding uncharacterized RLPs and RLKs were upregulated (**Fig. 6 K**). Most of these genes were not differentially expressed in RW. Some genes involved in response to wounding were upregulated in B3B5A and B3, such as two genes coding for homologs of linoleate 13S-lipoxygenases (**Fig. 6 I**). A gene encoding a phenylalanine ammonia-lyase (PAL) catalyzing the first step of the phenylpropanoid pathway was upregulated in RW, B3B5A and B3 (**Fig. 6 B**). Other induced genes in B3B5A and B3 participated in hypersensitive response. These included four genes coding for senescence-associated proteins (SAG12, SAG21, SAG101) and a membrane-attack complex/perforin (MACPF) domain-containing protein (**Fig. 6 B**), as well as a respiratory burst oxidase homolog protein F (RBOHF), a protein SRC2 mediating ROS production (**Fig. 6 E**), and a mitochondrial uncoupling protein contributing to the protection of plant cells against oxidative stress damage (**Fig. 6 E**). Defense signaling genes, and particularly calcium-mediated signaling genes, were also upregulated in B3B5A and B3 (**Fig. 6 C**). They comprised two genes coding for calcium-transporting ATPases, as well as genes encoding a calcium-binding protein and three glutamate receptors. Other upregulated genes involved in signaling encoded a lysine histidine transporter (**Fig. 6 C**), an enhanced disease susceptibility 1 (EDS1) and a protein Enhanced Disease Resistance 4 (EDR4), both essential for salicylic acid-mediated defense response, as well as two proteins NRT1/PTR involved in glucosinolate transport (**Fig. 6 B**). Once again, these genes were not differentially expressed in RW.

Some genes connected with the response to abiotic stimuli were also upregulated (**Fig. 6 G**). Genes encoding a CBS domain-containing protein and a putative E3 ubiquitin-protein ligase were upregulated in RW, B3B5A and B3 (**Fig. 6 G**), while genes encoding respectively a probable carboxylesterase playing a role during hypoxia, a TF involved in heat stress response (**Fig. 6 D**) and an aquaporin (**Fig. 6 G**) were only induced in B3B5A and B3.

In addition, in B3B5A and B3, 13 upregulated genes were involved in plant organ and tissue development, four of them being also induced in RW (**Fig. 6 L**). Six upregulated genes in B3B5A and B3, but not in RW, participated in sugar metabolism (**Fig. 6 J**). Eight genes were related to the response to light in relation to photoperiodism and circadian clock, six of them being also induced in RW (**Fig. 6 H**). Genes contributing to lipid and fatty acid metabolism were also upregulated, such as a gene encoding a probable Co-A ligase also induced in RW, and five genes encoding fatty acid desaturases, two being also found in RW (**Fig. 6 I**). In addition, a gene coding for a probable E3 ubiquitin ligase SUD1 involved in cuticle development was also upregulated in B3B5A and B3, but not in RW (**Fig. 6 I**). Also, a *BIM1* TF regulating brassinosteroid-induced genes was upregulated in B3B5A and B3 (**Fig. 6 D**). Three genes were associated with cell wall biogenesis or metabolism, they encoded a UDP-arabinose 4-epimerase, a probable pectinesterase inhibitor and a UDP-glucose 6-dehydrogenase, but only the first one was also induced in RW (**Fig. 6 F**).

Downregulated genes in RW, B3B5A and B3 included genes involved in plant organ development (**Fig. 6 L**), such as shoot and flower growth, as well as cell differentiation. Four genes related to the response to light were downregulated in B3B5A and B3, but only one was shared with RW (**Fig. 6 H**). Downregulated TFs in B3B5A and B3 comprised two heat-stress TFs, a *MYBS3* involved in cold tolerance and a basic leucine zipper contributing to cation homeostasis and response to salt stress (**Fig. 6 D**). In addition, a *NAC* TF acting as a positive regulator of age-dependent or stress-induced senescence and three genes coding for proteins OXIDATIVE STRESS 3 acting in chromatin remodeling upon stress were also downregulated in RW (**Fig. 6 D**). Two genes were associated with cell wall and lignin biogenesis (**Fig. 6 F**), and two other genes were part of the phosphatidylinositol metabolism (**Fig. 6 I**). Some genes involved in plant defenses were also downregulated in B3B5A and B3. It was the case for genes related to the response to oxidative stress, such as genes encoding respectively a peroxidase removing H_2_O_2_ upon pathogen attack and two thioredoxin-like proteins involved in cell redox homeostasis, a thioredoxin-like 1-2 being also downregulated in RW (**Fig. 6 E**). Additionally, a gene coding for a bifunctional nuclease participating in abscisic acid-derived callose deposition following pathogen infection was downregulated in B3B5A and B3 (**Fig. 6 B**). A cluster of three genes encoding 3beta,22alpha-dihydroxysteroid 3-dehydrogenases involved in brassinosteroid biosynthesis was repressed in RW, B3B5A and B3 (**Fig. 6 B**). These comparisons at the gene level are consistent with GO enrichment analyses conducted on single genotypes.

#### DEGs associated with QTL B5A

Eighteen DEGs were uniquely shared between B3B5A, B5AB.1 and B5AB.2 and were all upregulated, but four genes lacked a functional annotation (**Online Resource 6**). None of these genes were shared with RW (**Fig. 5 C**). Few genes displayed functions that could be directly related to defense response, except a gene encoding a probable disease resistance protein potentially involved in defense response to fungus (**Fig. 7A**). Other genes that could have a role in response to stress encoded respectively a putative UDP-glucose flavonoid 3-O-glucosyltransferase (**Fig. 7 A**), a 12-oxophytodienoate reductase contributing to jasmonic acid biosynthetic process (**Fig. 7 C**), a phenolic glucoside malonyltransferase associated with the response to toxic substance, and an AAA-ATPase ASD potentially implicated in responses to abiotic stresses (**Fig. 7 D**). A gene annotated as a Kunitz-type trypsin inhibitor (KPI106) involved in the establishment of symbiotic fungi was also expressed (**Fig. 7 I**). Two genes associated with cell walls and membranes were upregulated: they encoded a wall-associated receptor kinase contributing to the control of cell expansion (**Fig. 7 H**) and a protein DMP5 involved in membrane remodeling (**Fig. 7 E**). Other genes encoded two NAC domain-containing TFs and two U-box domain-containing proteins acting as E3 ubiquitin ligases (**Fig. 7 J**).

**Fig. 7.**
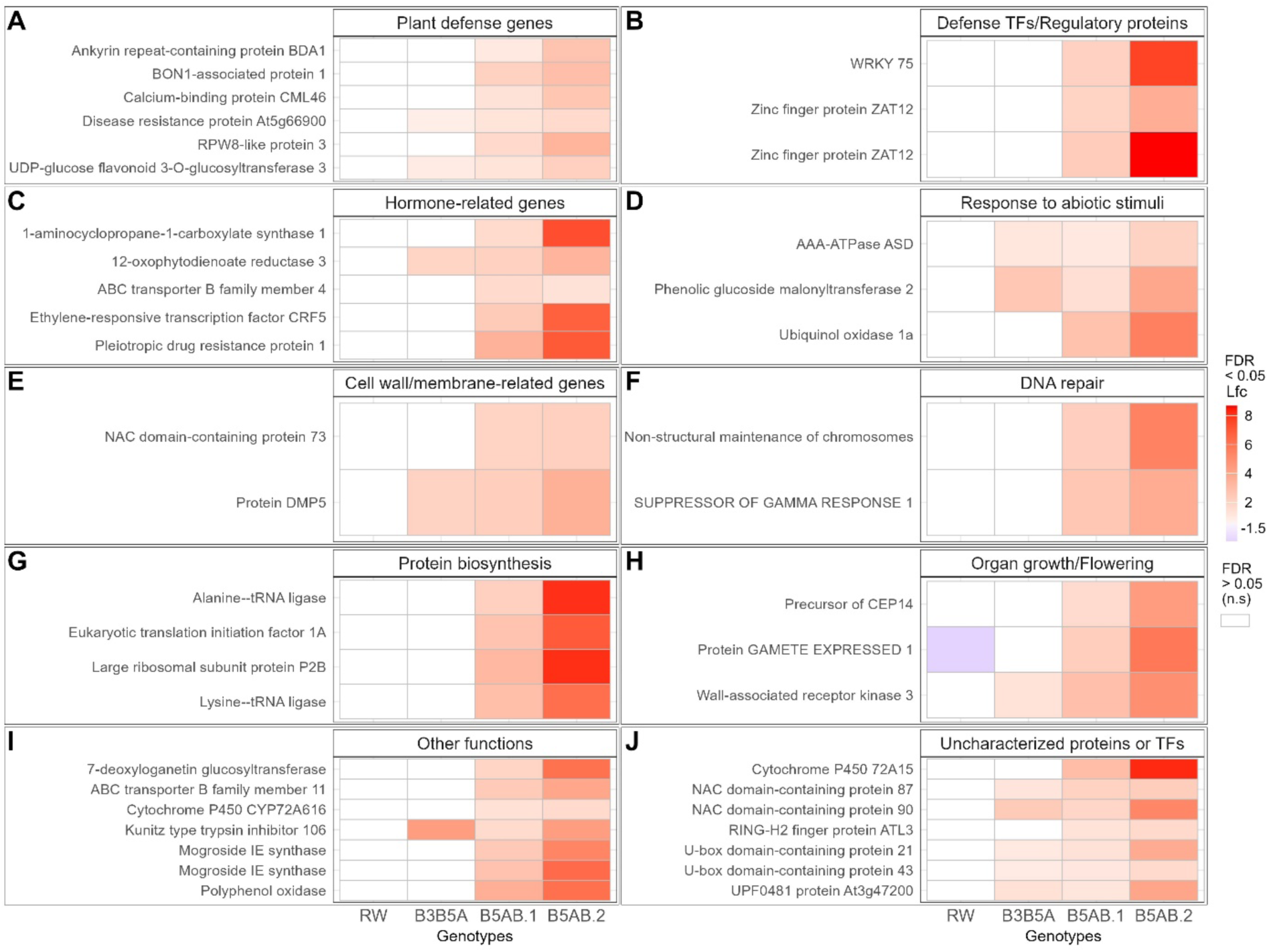
Heatmaps of annotated common DEGs between RW, B3B5A, B5AB.1 and B5AB.2 at 3 dpi. DEGs were classified according to their associated keywords or gene families. Genes not significantly (n.s.) differentially expressed are displayed in white. Fold changes, compared to the non-inoculated leaves, of down- (blue) and upregulation (red) are shown on a log2 scale (lfc). (A) Plant defense genes; (B) Defense TFs/Regulatory proteins; (C) Hormone-related genes; (D) Response to abiotic stimuli; (E) Cell wall/membrane-related genes; (F) DNA repair; (G) Protein biosynthesis; (H) Organ growth/ Flowering; (I) Other functions; (J) Uncharacterized proteins or TFs

#### DEGs associated with QTL B5B

Thirty-four genes were uniquely shared between B5AB.1 and B5AB.2 and were all upregulated. Six were not or poorly annotated and are not displayed on Fig. 7 (**Online Resource 6**). These common DEGs comprised genes involved in defense response, such as genes coding for a probable WRKY75 and two homologs of a zinc finger protein ZAT12, all three genes being TFs regulating the response to fungi (**Fig. 7 B**). A gene encoding a negative regulator of cell death and plant defenses, BON1-associated protein 1 (BAP1), was also upregulated, as well as a *RESISTANCE TO POWDERY MILDEW 8 (RPW8)-like 3* (**Fig. 7 A**). Concerning defense signaling, gene coding for an ankyrin repeat-containing protein BDA1 and one coding for a probable calcium-binding protein CML46 (**Fig. 7 A**) were upregulated. Moreover, genes encoding a 1-aminocyclopropane-1-carboxylate synthase 1 involved in ethylene biosynthesis and an ethylene-responsive transcription factor CRF5 were also induced, as well as one encoding a pleiotropic drug resistance protein 1 involved in strigolactone transport and an auxin transporter ABC transporter B family member 4 (**Fig. 7 C**). A NAC domain-containing TF regulating cell wall biogenesis was also upregulated (**Fig. 7 E**). Two genes involved in DNA repair (**Fig. 7 F**) and four genes playing roles in protein biosynthesis were induced (**Fig. 7 G**). A gene encoding the protein GAMETE EXPRESSED 1 was also found in RW, but was downregulated, unlike in B5AB.1 and B5AB.2 (**Fig. 7 H**).

## Discussion

The present study aimed to identify the defense mechanisms associated with major QTLs for black spot resistance derived from *R. wichurana*. To this end, we compared the transcriptomic responses of five genotypes, each harboring different QTLs or QTL combinations, upon interaction with *D. rosae*.

### Ups and Downs: a singular regulation of defense-related genes in RW

In addition to its relatively small number of DEGs, unusual patterns of defense gene expression were displayed at 3 dpi in RW. Indeed, few genes involved in pathogen detection, defense signaling or regulation of transcription were upregulated, and the majority were downregulated, which is quite surprising. As we are dealing with a resistant genotype, we would expect defense genes to be overexpressed or not differentially expressed, as observed in other studies for the interaction rose-*D. rosae* (Song et al. 2024) and for other pathosystems (e.g. McGregor et al., 2009; Dmitriev et al., 2017; Chandran et al., 2021; Su et al., 2023; Ren et al., 2024).

In brief, RW possessed a different expression pattern compared to the F1 genotypes carrying QTLs B5A and B5B, but some similarities with F1 genotypes carrying QTL B3. Contrary to its progenies, RW harbors its entire genetic background, containing at least two minor QTLs on chromosomes 4 and 6 (Lopez Arias et al. 2020). This genetic background in combination with the QTLs B3 and B5 may contribute to an earlier and/or more efficient defense response of RW. The phenotypic data, the GO analysis as well as the observation of the DEGs corroborate this hypothesis. Indeed, the SEA of GO terms showed that many processes that were upregulated in the F1 genotypes at 3 dpi were downregulated in RW (**Fig. 4, 6** and **7**). This might indicate the end of the defense response in RW and a return to normal expression, with some genes being no more differentially expressed, and some processes being “forced back” to normal after accumulation of the proteins and metabolites they produced during infection. In this case, 3 dpi may have been too late to be able to compare the reactions between RW and its progenies. Considering that RW’s resistance is quantitative, it is likely that each F1 genotype inherited some minor-effect QTLs that have not been detected yet but that participate in its resistance phenotype. The effect of each genotype’s genetic background should not be ignored and the originality concerning the plant material used (F1 progenies, not Near Isogenic Lines) should be taken into account when examining the current RNA-seq analysis.

### GO terms associated with the infection by *D. rosae*

The apparition of symptoms and the transcriptomic changes in the five rose genotypes confirmed their responses to *D. rosae* inoculation. GO enrichment and examination of DEGs identified terms and genes associated with response to biotic stimuli in general and with defense response to fungus in particular, except in RW. Transcriptome analyses performed by Neu et al. (2019) and Song et al. (2024) on susceptible rose genotypes showed that terms such as “response to biotic stimulus”, “defense response”, and “chitin catabolic process” were enriched. However, Song et al. (2024) did not find these terms when studying a resistant genotype and hypothesized that these terms may be common to susceptible roses infected by *D. rosae*. In the present work, the four F1 genotypes were partially resistant to black spot (**Fig. 2**) and exhibited enrichment in similar terms (**Fig. 4** and **Online Resource 5**). In contrast, terms related to defense were not found in the resistant parent RW, and the term “response to stress” was associated with downregulated genes, unlike in the four F1 genotypes. Genes related to membranes were upregulated in B3B5A, B3 and B5AB.2, suggesting that these genotypes may utilize membrane-associated processes in their defense. Indeed, many receptors specialized in pathogen recognition are located in the plasma membrane, and transmembrane transport is key in plant immune system (Ngou et al. 2022).

#### Detection of *D. rosae*, QTL specificity

Many genes encoding receptors and disease resistance proteins were identified among the five genotypes. RLPs, RLKs or WAKs are transmembrane receptors and constitute the first layer in plant defense as their extracellular domains enable the early detection of pathogens (Ngou et al. 2022). Among the common DEGs between B3B5A and B3, eight genes were annotated as RLPs or RLKs related to defense (**Fig. 6 A**), and eight other genes were annotated as RLPs or RLKs but their precise biological function was unknown (**Fig. 6 K**). RLPs play key roles in plant growth and development, but also in various stress responses, including pathogen attack (Wang et al. 2008). Therefore, we can hypothesize that some of the uncharacterized RLPs transcripts accumulated here could be involved in *D. rosae* recognition and be controlled by QTL B3. Among these RLPs, only one proline-rich receptor-like protein kinase PERK1 responding to wounding and fungal infestation (Silva and Goring 2002) was also upregulated in RW. Defense responses also require regulators of various protein natures. In B5AB.1 and B5AB.2, the gene *BDA1* was upregulated (**Fig. 7 A**). This protein functions as an essential signaling component downstream of the RLP SNC2, a key regulator of defense responses (Yang et al. 2012). In these genotypes, a gene encoding a BAP1 was also upregulated. This small protein and its partner protein BON1 negatively regulate the salicylic-acid mediated cell death through inhibition of *R* gene *SNC1* in Arabidopsis (Yang et al. 2006; Yang et al. 2007).

The second layer of defense relies on intracellular nucleotide-binding leucine-rich repeat receptors (NLRs) (Ngou et al. 2022). NLRs appeared to be very diverse in our datasets. Numerous NLRs were commonly upregulated in B3B5A and B3 (**Fig. 6 A**). For instance, two homologs of a gene encoding a protein Resistance to *Uncinula necator* (RUN1) were upregulated in both genotypes. *RUN1* belongs to the *TNL* genes (a subclass of NLRs), and confers resistance to powdery mildew in grapevine (Feechan et al. 2013). Two other *TNLs*, encoding Tobacco Mosaic Virus (TMV) resistance protein N homologs, were also upregulated in B3B5A and B3 in the present study, and in RW in the study of Lopez Arias (2021).

In addition, an *MLO-like protein 1*, an *MLO-like protein 6* and a *DOWNY MILDEW RESISTANCE 6* (*DMR6*) were upregulated in B3B5A and B3. Remarkably, they are considered as susceptibility genes (Zeilmaker et al. 2015; Kusch and Panstruga 2017). *MLO-like* genes provide partial resistance to different species of powdery mildew in various plants by inducing rapid callose apposition at the points of fungal penetration, possibly through efficient calcium signaling (Jørgensen 1992; Kusch and Panstruga 2017). *MLO-like* genes were also induced during defense response to other pathogens, such as rusts (*Uromyces* spp.) in grass pea (Almeida et al. 2014) and six pathogens of apple, including the bacteria *Erwinia amylovora*, the hemibiotrophic fungi *Venturia inaequalis* and *Diplocarpon mali*, the necrotropic fungi *Valsa mali* and *Alternaria alternate*, and the necrotrophic oomycete *Pythium ultimum* (Balan et al. 2018). Interestingly, another powdery mildew resistance-like gene was upregulated in B5AB.1 and B5AB.2 (**Fig. 7 A**), namely *RPW8-like protein 3*. In Arabidopsis, RPW8 conferred resistance to a broad spectrum of powdery mildew strains, as well as to cauliflower mosaic virus, and to oomycetes such as *Hyaloperonospora parasitica* (Wang et al. 2007) and *H. arabidopsidis* (Ma et al. 2014). *RPW8* gene family actually acts as a regulator of salicylic acid-dependent basal defenses, rather than as powdery mildew-specific *R* genes (Li et al. 2020). This hypothesis would be consistent with our results in the rose-*D. rosae* pathosystem. Regarding B3B5A, B5AB.1 and B5AB.2, transcripts encoding a homolog of the coiled-coil NLR (CNL) At5g66900 were accumulated. This CNL acted as a helper TNL in reaction to the oomycete *H. arabidopsidis* and the bacterium *P. syringae* (Wu et al. 2019). Helper NLRs operate downstream of classic NLRs: when classic NLRs detect a pathogenic attack, they activate the helper NLRs, which amplify and propagate the defense signal throughout the cell, contributing to an effective, coordinated immune response.

### ROS production and callose deposition

Recently, plant defense mechanisms have been shown to be widely intertwined in space and time, and not as sequential as implied by the initial zigzag model by Jones and Dangl (2006). The joint activation of extracellular receptors and *NLR* genes results in ROS accumulation, callose apposition, and further defense mechanisms (Ngou et al. 2022).

In B3B5A and B3, genes associated with ROS production were upregulated while genes involved in the protection of plant cells against oxidative damage were both up- and downregulated, pointing to a fine-tuning of ROS homeostasis (**Fig. 6 E**). Indeed, ROS accumulation can lead to cell death (Andersen et al. 2018; Ngou et al. 2022), but at moderate concentrations, they contribute to the modulation of defense response (Kaur et al. 2022). For example, a gene coding for the respiratory burst oxidase RBOHF was upregulated in B3B5A and B3. RBOHF was shown to be an important regulator of defense in Arabidopsis (Arnaud et al. 2023).

ROS accumulation also leads to callose deposition (Ngou et al. 2022). Callose deposition has been frequently observed in the rose-*D. rosae* interaction (Gachomo 2005; Blechert and Debener 2005; Lopez Arias 2021; Yang et al. 2022), but transcriptomic studies did not reveal genes such as callose synthases , whereas they are thought to play important roles in callose- mediated defenses in plants (Voigt and Somerville 2009). In B3B5A, B3 and B5AB.2, the term “regulation of defense response by callose deposition” was upregulated (**Fig. 4 C**), but no common DEGs between B3B5A and B3, nor between B3B5A, B5AB.1 and B5AB.2 were actually related to callose biosynthesis. This suggests that different DEGs were involved in callose deposition in these genotypes. Callose also acts in response to wounding by strengthening cell walls and maintaining cell integrity (Voigt and Somerville 2009).

More generally, cell wall modifications play critical roles in plant defenses (Andersen et al. 2018). In B3B5A and B3, genes associated with cell wall biogenesis and lignin biosynthesis were both up- and downregulated, suggesting a nuanced control of these processes (**Fig. 6 F**). In B5AB.1 and B5AB.2, a NAC domain-containing protein 73 involved in cell wall biogenesis was also induced (**Fig. 7 E**). This TF regulates cellulose and hemicellulose biosynthesis as well as lignin polymerization and signaling (Hussey et al. 2011). In rose, cell wall modifications have been reported in the case of susceptibility to *P. pannosa*, powdery mildew causal agent (Neu et al. 2019), but different genes were involved. In addition, controlled cell death appeared to be a significant part of defense, particularly in B3B5A and B3 (**Fig. 4 C**). However, since these genotypes are only partially resistant, this response is not completely successful at stopping the fungus.

### Transcriptional and translational regulations upon infection

Regarding defense-related TFs, a *WRKY40* was induced in B3B5A and B3 (**Fig. 6 D**), which is consistent with previous results in rose. Indeed, in both resistant and susceptible rose genotypes infected by *D. rosae*, WRKY40 acted as a regulator of the antagonistic salicylic-jasmonic acid pathway (Zheng et al. 2024). WRKY40 was also associated with aphid resistance in rose (Dong et al. 2024). In B5AB.1 and B5AB.2, a *WRKY75* was upregulated (**Fig. 7 B**). This TF was also expressed in the study of Domes and Debener (2024) in the rose-*D.rosae* pathosystem. Interestingly, it was upregulated at 24 and 72 hpi in the resistant genotype, but only at 24 hpi in the susceptible one. Unlike what was found in Arabidopsis, the homologs of *WRKY75* in rose and strawberry did not act as initiators of phytohormone-dependent immune responses (Domes and Debener 2024 and references therein). Two genes coding for a homolog of the zinc-finger protein ZAT12 were also upregulated in B5AB.1 and B5AB.2. This TF was mostly studied in Arabidopsis and was found in stress conditions such as wounding, infection by bacteria, fungi and nematodes or contact with their elicitors, but also oxidative stress, heat, cold, water deprivation and UV application (Davletova et al. 2005).

In addition to TFs, genes involved in protein translation were upregulated in B5AB.1 and B5AB.2 (**Fig. 7 G**), suggesting that the response to *D. rosae* through protein generation was still ongoing at 3 dpi.

### Potential other defense mechanisms

Due to their antimicrobial properties, phenylpropanoids and alkaloids are part of phytoalexins, which are toxic metabolites involved in plant defense response against pathogens (Andersen et al. 2018). In B3B5A and B3, two DEGs were associated with phenylpropanoid pathways, they encoded a PAL (also expressed in RW) and a regulatory protein homolog of At2g44130 (**Fig. 6 B, D**). In B3B5A, B5AB.1 and B5AB.2, a gene coding for a UDP-glucose flavonoid 3-O-glucosyltransferase was also expressed. In the studies conducted by Neu et al. (2019) and Song et al. (2024), GO terms related to phenylpropanoid and flavonoid pathways were also enriched in upregulated genes, indicating that these compounds could be important for the resistance of rose to *D. rosae*.

Interestingly, B3B5A, B5AB.1 and B5AB.2 overexpressed a homolog of a gene encoding the Kunitz protease inhibitor KPI106 (**Fig. 7 I**), which was also found in RW in a previous study (Lopez Arias 2021). This secreted protease inhibitor was described in *Medicago truncatula* as controlling mycorrhiza development during root colonization by symbiotic fungi (Rech et al. 2013). The authors suggested that, during the interaction with a symbiont, a protease detected an unknown target and generated a signaling peptide enabling fungal development within the root. KPI106 would modulate the activity of the protease by competing with its substrate. In our case, KPI106 was expressed upon pathogen infection. Many protease inhibitors and Kunitz protease inhibitors are involved in plant defense against pathogens and pests (Rech et al. 2013; Arnaiz et al. 2018 and references therein). Therefore, we can hypothesize that this KPI106 homolog possesses antifungal properties, thus protecting plant cells against *D. rosae*.

### Connections between stress response mechanisms

The analysis also yielded terms and genes that are usually associated with the response to abiotic factors, such as light, cold, salinity, hypoxia and nitrogen compounds. This suggests that common regulations exist between biotic and abiotic stimuli in rose.

The GO term “response to light stimulus” was globally downregulated in RW, B3B5A, and B3. However, common genes between these three genotypes were both upregulated and downregulated (**Fig. 6 H**), e.g. genes responsive to UV-A, UV-B, red light and blue light. Additionally, in B3B5A and B3, five genes involved in photoperiodism and circadian rhythm were differentially expressed, four being upregulated and shared with RW. Recent studies have unveiled the major role of clock genes in plant defense mechanisms (Lu et al. 2017; Galeou et al. 2023; Ren et al. 2024).

Concerning response to temperature, both responses to heat and cold were differentially regulated in the genotypes studied here. In B3B5A, B3 and B5AB.2, several genes encoding heat shock proteins and heat stress TFs were differentially expressed (**Fig. 6 D** and **Online Resource 6**), similar to another study on the rose-*D. rosae* pathosystem (Neu et al. 2019). These genes, also called heat shock factors (HSFs), are transcription regulators of various stress-inducible genes, such as chaperones (heat shock proteins and others), other TFs, but also *PR* genes and genes involved in hormonal pathways (Andrási et al. 2021). During pathogen infestation, HSFs modulate ROS signaling, oxidative stress and cell death, thus constituting important components of disease resistance in plants (Hoang et al. 2019; Andrási et al. 2021). B3B5A and B3 exhibited four upregulated low temperature-responsive genes and one downregulated one. Among these genes, *SRC2* (**Fig. 6 E**) and *SAG21* (**Fig. 6 B**) also participated in the defense response. Interestingly, in Arabidopsis, SRC2 activated calcium-mediated ROS production by RBOHF (Kawarazaki et al. 2013), which was also expressed in B3B5A and B3 (**Fig. 6 E**). As for *SAG21*, it constitutes a great example of hub gene for diverse physiological processes, as it is involved in growth regulation and in response to various stimuli: light, cold, drought, salinity, wounding, *Botrytis cinerea* infection and both fungal and bacterial elicitors (Evans et al. 2024).

### Crosstalk between actors of signaling

The activation or repression of hormone signal transduction pathways upon *D. rosae* infection appears to be genotype-dependent in roses, affecting both the nature of the hormones involved and the intensity of the response (Neu et al. 2019; Song et al. 2024). In our study, genes playing roles in salicylic acid biosynthesis, such as *EDS1* and *SAG101*, were upregulated in B3B5A and B3 (**Fig. 6 B**), but they were not differentially regulated in RW. These two genes were detected in the rose-*D. rosae* pathosystem by Neu et al. (2019), but not by Song et al. (2024). Ethylene acts as an important defense mediator in various plants including rose. Neu et al. (2019) reported a contradictory expression of ethylene-related genes in their susceptible genotype infected by *D. rosae*. As for Song et al. (2024), they only found two homologs of *ERFs* upregulated in their susceptible genotype, while no ethylene-related genes were upregulated for the resistant one. In our study, an ethylene-responsive TF was upregulated in B5AB.1 and B5AB.2 (**Fig. 7 C**). Abscisic acid (ABA) signaling was globally downregulated in RW and B3, and upregulated in B3B5A and B5AB.2 (**Fig. 4 E**). B3B5A and B3 both underexpressed three genes of the *OXIDATIVE STRESS 3* family (**Fig. 6 D**), which have roles in chromatin remodeling and act as negative regulators of several ABA-responsive genes (Xiao et al. 2021). In the rose-*D. rosae* pathosystem, infected rose leaves produced ABA, unlike non-infected leaves. Interestingly, ABA was also produced by *D. rosae* grown in potato dextrose medium, possibly indicating that ABA plays a role in chlorosis and leaf abscission in compatible interactions (Debener 2019). In the resistant rose studied by Song et al. (2024), ABA response seemed to vary depending on the time after inoculation and on the studied gene in ABA signaling pathway. Genes involved in the biosynthesis of brassinosteroids were downregulated in RW, B3B5A and B3 (**Fig. 6 B**). Depending on the pathosystem, brassinosteroids have been reported to positively or negatively regulate defense (Yu et al. 2018). On a susceptible rose, Song et al. (2024) brought to light a potential negative effect of brassinosteroids on black spot resistance. Therefore, the repressions of brassinosteroid biosynthesis in the resistant RW and partially resistant B3B5A and B3 are consistent with their finding.

GO terms associated with calcium signaling were significantly enriched in the four F1 genotypes (**Fig. 4 E**, **Fig. 6 C**, **Fig. 7 A**). Calcium signaling plays a crucial role in plant defenses by acting as a second messenger in response to various biotic stresses. It controls the activation of hormones, defense-related genes, as well as the production of ROS, which are essential for initiating further defense response (Bhar et al. 2023). Indeed, calcium-binding proteins regulate genes such as *EDS1*, and cellular calcium levels also trigger *RBOHs* (Bhar et al. 2023), and *MLO*-like genes (Kusch and Panstruga 2017), these three genes being overexpressed in B3B5A and B3.

## Conclusions

Despite some limitations, the present study disclosed a wide range of defense responses of roses derived from *R. wichurana* to black spot disease, and pointed out the considerable intricacy of these processes.

Based on these data, relying on a few individuals, it is not easy to precisely define the roles of QTLs B3, B5A and B5B inherited from RW, especially since GO terms were shared between genotypes of different QTL classes and we may have lacked statistical power for some genotypes. However, we can make some assumptions, particularly for QTL B3 for which the two representative genotypes shared numerous DEGs (**Fig. 8**).

**Fig. 8.**
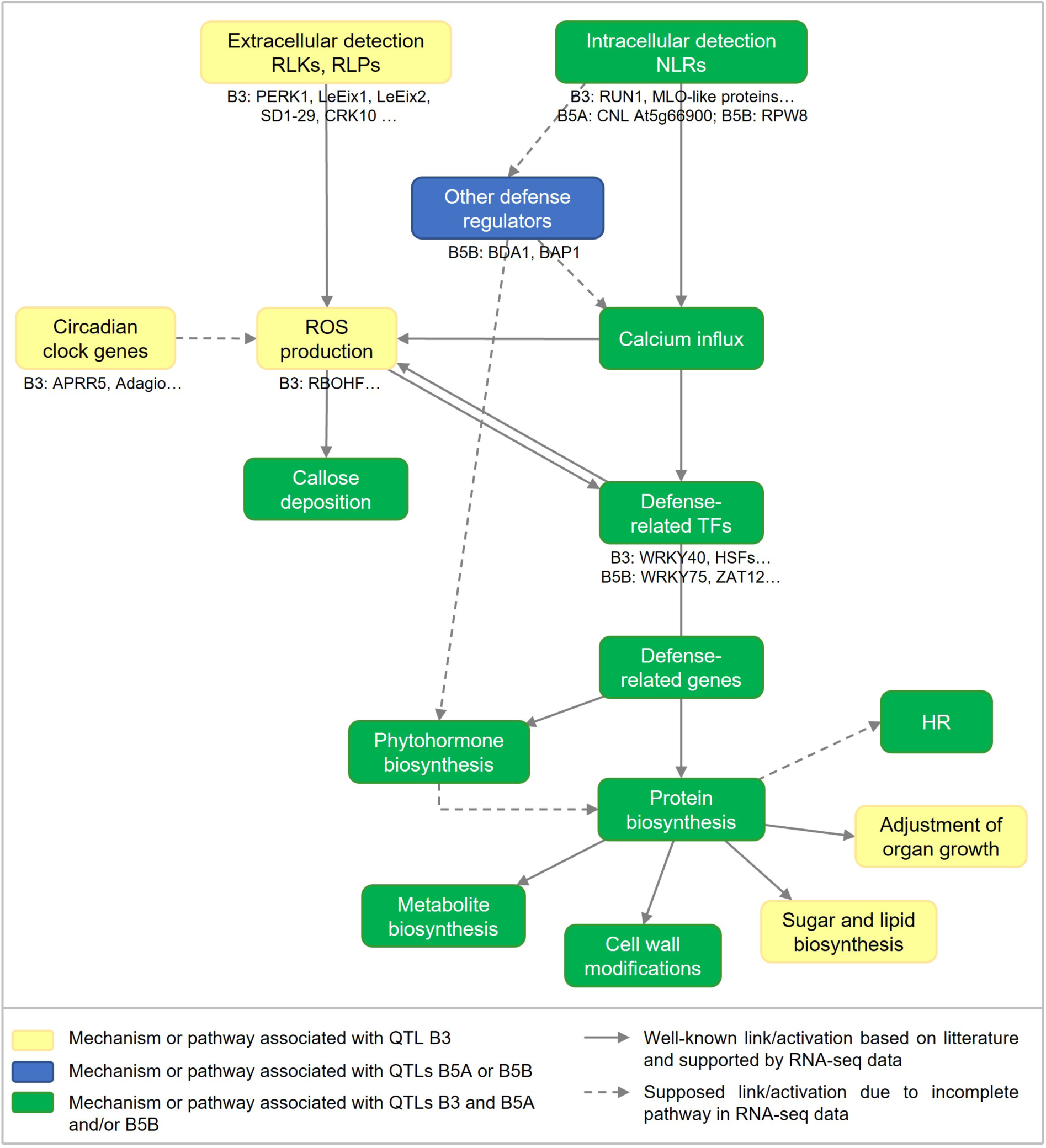
Components of the response of RW to black spot disease, based on transcriptomic data from RW and four F1 progenies harboring different QTL combinations

Genotypes sharing QTL B3 expressed “classic” defense mechanisms. Indeed, it involved pathogen detection through extracellular receptors and NLRs, followed by defense signaling cascades including hormones (particularly salicylic and abscisic acids), calcium, lipid and sugar signaling. All these processes led to ROS production, callose apposition and probable localized cell death. Many genes that have been discussed here constitute interesting candidates in the global defense response associated with this QTL. Conversely, genotypes harboring QTL B5 alone did not exhibit many DEGs, and shared even fewer. The low number of common DEGs makes it difficult to connect them with common mechanisms that could be proper to the QTLs on chromosome B5, although some interesting mechanisms have been revealed.

## Supporting information

Online Resource 1

Online Resource 2

Online Resource 3

Online Resource 4

Online Resource 5

Online Resource 6

## Statements & Declarations

### Author Contributions

FF, LHSO, SPa and VSF conceived the study. CB, CV, JC and TT participated in material preparation. JC, JJ, LHSO, LL, SPa and TT sampled and phenotyped the plants. JJ carried out RNA extraction and RNA quality control. LL performed transcriptomic data analyses. DCLA, JJ, SA, SPa and SPe provided support in bioinformatic and transcriptomic analyses. LL wrote the manuscript. FF, SPa and VSF provided extensive revisions of the manuscript. All authors read and approved the final manuscript.

### Data Availability

Raw transcriptomic data are available on the European Nucleotide Archive website under bioproject PRJEB90172.

## Acknowledgments

We are grateful for ANAN platform collaboration for DNA extraction. We thank Phenotic platform (https://doi.org/10.17180/YKBZ-2V85) and INRAE Horticulture Experimental Facility (Beaucouzé, France) for plant management in greenhouse and experimental fields, respectively.

## References

Almeida NF, Leitão ST, Krezdorn N, Rotter B, Winter P, Rubiales D, Vaz Patto MC (2014) Allelic diversity in the transcriptomes of contrasting rust-infected genotypes of Lathyrus sativus, a lasting resource for smart breeding. BMC Plant Biol 14:376. 10.1186/s12870-014-0376-2

Andersen E, Ali S, Byamukama E, Yen Y, Nepal M (2018) Disease Resistance Mechanisms in Plants. Genes 9:339. 10.3390/genes9070339

Andrási N, Pettkó-Szandtner A, Szabados L (2021) Diversity of plant heat shock factors: regulation, interactions, and functions. Journal of Experimental Botany 72:1558–1575. 10.1093/jxb/eraa576

Arnaiz A, Talavera-Mateo L, Gonzalez-Melendi P, Martinez M, Diaz I, Santamaria ME (2018) Arabidopsis Kunitz Trypsin Inhibitors in Defense Against Spider Mites. Front Plant Sci 9. 10.3389/fpls.2018.00986

Arnaud D, Deeks MJ, Smirnoff N (2023) RBOHF activates stomatal immunity by modulating both reactive oxygen species and apoplastic pH dynamics in Arabidopsis. The Plant Journal 116:404–415. 10.1111/tpj.16380

Balan B, Marra FP, Caruso T, Martinelli F (2018) Transcriptomic responses to biotic stresses in Malus x domestica: a meta-analysis study. Sci Rep 8:1970. 10.1038/s41598-018-19348-4

Benjamini Y, Hochberg Y (1995) Controlling the False Discovery Rate: A Practical and Powerful Approach to Multiple Testing. Journal of the Royal Statistical Society: Series B (Methodological) 57:289–300. 10.1111/j.2517-6161.1995.tb02031.x

Benjamini Y, Yekutieli D (2001) The control of the false discovery rate in multiple testing under dependency. The Annals of Statistics 29:1165–1188. 10.1214/aos/1013699998

Bhar A, Chakraborty A, Roy A (2023) The captivating role of calcium in plant-microbe interaction. Front Plant Sci 14. 10.3389/fpls.2023.1138252

Blechert O, Debener T (2005) Morphological characterization of the interaction between Diplocarpon rosae and various rose species. Plant Pathology 54:82–90. 10.1111/j.1365-3059.2005.01118.x

Byrne DH, Anderson N, Brent Pemberton H (2007) The Use of Rosa wichurana in the Development of Landscape Roses Adapted to Hot Humid Climates. Acta Hortic 267– 274. 10.17660/ActaHortic.2007.751.34

Chandran NK, Sriram S, Prakash T, Budhwar R (2021) Transcriptome changes in resistant and susceptible rose in response to powdery mildew. Journal of Phytopathology 169:556–569. 10.1111/jph.13028

Chen S, Zhou Y, Chen Y, Gu J (2018) fastp: an ultra-fast all-in-one FASTQ preprocessor. Bioinformatics 34:i884–i890. 10.1093/bioinformatics/bty560

Davletova S, Schlauch K, Coutu J, Mittler R (2005) The Zinc-Finger Protein Zat12 Plays a Central Role in Reactive Oxygen and Abiotic Stress Signaling in Arabidopsis. Plant Physiology 139:847–856. 10.1104/pp.105.068254

Debener T (2019) The Beast and the Beauty: What Do we know about Black Spot in Roses? Critical Reviews in Plant Sciences 38:313–326. 10.1080/07352689.2019.1665778

Dmitriev AA, Krasnov GS, Rozhmina TA, Novakovskiy RO, Snezhkina AV, Fedorova MS, Yurkevich OYu, Muravenko OV, Bolsheva NL, Kudryavtseva AV, Melnikova NV (2017) Differential gene expression in response to Fusarium oxysporum infection in resistant and susceptible genotypes of flax (Linum usitatissimum L.). BMC Plant Biol 17:253. 10.1186/s12870-017-1192-2

Domes HS, Debener T (2024) Genome-Wide Analysis of the WRKY Transcription Factor Family in Roses and Their Putative Role in Defence Signalling in the Rose–Blackspot Interaction. Plants 13:1066. 10.3390/plants13081066

Dong W, Sun L, Jiao B, Zhao P, Ma C, Gao J, Zhou S (2024) Evaluation of aphid resistance on different rose cultivars and transcriptome analysis in response to aphid infestation. BMC Genomics 25:232. 10.1186/s12864-024-10100-z

Du Z, Zhou X, Ling Y, Zhang Z, Su Z (2010) agriGO: a GO analysis toolkit for the agricultural community. Nucleic Acids Research 38:W64–W70. 10.1093/nar/gkq310

Evans KV, Ransom E, Nayakoti S, Wilding B, Mohd Salleh F, Gržina I, Erber L, Tse C, Hill C, Polanski K, Holland A, Bukhat S, Herbert RJ, de Graaf BHJ, Denby K, Buchanan-Wollaston V, Rogers HJ (2024) Expression of the Arabidopsis redox-related LEA protein, SAG21 is regulated by ERF, NAC and WRKY transcription factors. Sci Rep 14:7756. 10.1038/s41598-024-58161-0

Ewels P, Magnusson M, Lundin S, Käller M (2016) MultiQC: summarize analysis results for multiple tools and samples in a single report. Bioinformatics 32:3047–3048. 10.1093/bioinformatics/btw354

Feechan A, Anderson C, Torregrosa L, Jermakow A, Mestre P, Wiedemann-Merdinoglu S, Merdinoglu D, Walker AR, Cadle-Davidson L, Reisch B, Aubourg S, Bentahar N, Shrestha B, Bouquet A, Adam-Blondon A-F, Thomas MR, Dry IB (2013) Genetic dissection of a TIR-NB-LRR locus from the wild North American grapevine species Muscadinia rotundifolia identifies paralogous genes conferring resistance to major fungal and oomycete pathogens in cultivated grapevine. The Plant Journal 76:661–674. 10.1111/tpj.12327

Gachomo EW (2005) Studies of the life cycle of Diplocarpon rosae Wolf on roses and the effectiveness of fungicides on pathogenesis. PhD thesis, University of Bonn

Gachomo EW, Kotchoni SO (2007) Detailed description of developmental growth stages of Diplocarpon rosae in Rosa: a core building block for efficient disease management. Annals of Applied Biology 151:233–243. 10.1111/j.1744-7348.2007.00167.x

Galeou A, Stefanatou C, Prombona A (2023) Circadian clock-dependent and -independent response of Phaseolus vulgaris to Pseudomonas syringae. Physiological and Molecular Plant Pathology 124:101944. 10.1016/j.pmpp.2022.101944

Hattendorf A, Linde M, Mattiesch L, Debener T, Kaufmann H (2004) Genetic Analysis of Rose Resistance Genes and their Localisation in the Rose Genome. Acta Hortic 123– 130. 10.17660/ActaHortic.2004.651.14

Hoang TV, Vo KTX, Rahman MM, Choi S-H, Jeon J-S (2019) Heat stress transcription factor OsSPL7 plays a critical role in reactive oxygen species balance and stress responses in rice. Plant Science 289:110273. 10.1016/j.plantsci.2019.110273

Hussey SG, Mizrachi E, Spokevicius AV, Bossinger G, Berger DK, Myburg AA (2011) SND2, a NAC transcription factor gene, regulates genes involved in secondary cell wall development in Arabidopsis fibres and increases fibre cell area in Eucalyptus. BMC Plant Biology 11:173. 10.1186/1471-2229-11-173

Jørgensen IH (1992) Discovery, characterization and exploitation of Mlo powdery mildew resistance in barley. Euphytica 63:141–152. 10.1007/BF00023919

Kaur S, Samota MK, Choudhary M, Choudhary M, Pandey AK, Sharma A, Thakur J (2022) How do plants defend themselves against pathogens-Biochemical mechanisms and genetic interventions. Physiol Mol Biol Plants 28:485–504. 10.1007/s12298-022-01146-y

Kawarazaki T, Kimura S, Iizuka A, Hanamata S, Nibori H, Michikawa M, Imai A, Abe M, Kaya H, Kuchitsu K (2013) A low temperature-inducible protein AtSRC2 enhances the ROS-producing activity of NADPH oxidase AtRbohF. Biochim Biophys Acta 1833:2775–2780. 10.1016/j.bbamcr.2013.06.024

Kusch S, Panstruga R (2017) mlo-Based Resistance: An Apparently Universal “Weapon” to Defeat Powdery Mildew Disease. MPMI 30:179–189. 10.1094/MPMI-12-16-0255-CR

Labbé J (2014) LOI n° 2014-110 du 6 février 2014 visant à mieux encadrer l’utilisation des produits phytosanitaires sur le territoire national

Lau J, Gill H, Taniguti CH, Young EL, Klein PE, Byrne DH, Riera-Lizarazu O (2023) QTL discovery for resistance to black spot and cercospora leaf spot, and defoliation in two interconnected F1 bi-parental tetraploid garden rose populations. Front Plant Sci 14:1209445. 10.3389/fpls.2023.1209445

Leus L (2017) Selection Strategies for Disease Resistance in Roses. In: Reference Module in Life Sciences. Elsevier, p B9780128096338051000

Li L, Habring A, Wang K, Weigel D (2020) Atypical Resistance Protein RPW8/HR Triggers Oligomerization of the NLR Immune Receptor RPP7 and Autoimmunity. Cell Host & Microbe 27:405–417.e6. 10.1016/j.chom.2020.01.012

Lopez Arias DC (2021) Genetics and genomics of black spot disease resistance in garden roses. PhD thesis, Agrocampus Ouest

Lopez Arias DC, Chastellier A, Thouroude T, Bradeen J, Van Eck L, De Oliveira Y, Paillard S, Foucher F, Hibrand-Saint Oyant L, Soufflet-Freslon V (2020) Characterization of black spot resistance in diploid roses with QTL detection, meta-analysis and candidate-gene identification. Theor Appl Genet 133:3299–3321. 10.1007/s00122-020-03670-5

Lopez Arias DC, Zlesak DC, Zurn JD, Moore EM, Clark M, Bassil NV, Hokanson SC (2023) Rdr5, a Novel Rose Black Spot Disease Resistance Locus in the Tetraploid Garden Rose Ramblin’Red

Love MI, Huber W, Anders S (2014) Moderated estimation of fold change and dispersion for RNA-seq data with DESeq2. Genome Biol 15:550. 10.1186/s13059-014-0550-8

Lu H, McClung CR, Zhang C (2017) Tick Tock: Circadian Regulation of Plant Innate Immunity. Annual Review of Phytopathology 55:287–311. 10.1146/annurev-phyto-080516-035451

Ma X-F, Li Y, Sun J-L, Wang T-T, Fan J, Lei Y, Huang Y-Y, Xu Y-J, Zhao J-Q, Xiao S, Wang W-M (2014) Ectopic Expression of RESISTANCE TO POWDERY MILDEW8.1 Confers Resistance to Fungal and Oomycete Pathogens in Arabidopsis. Plant and Cell Physiology 55:1484–1496. 10.1093/pcp/pcu080

Marolleau B, Petiteau A, Bellanger M, Sannier M, Le Pocreau N, Porcher L, Paillard S, Foucher F, Thouroude T, Serres-Giardi L, Aguileta G, Chastellier A, Bonneau C, Le Cam B, Soufflet-Freslon V, Hibrand-Saint Oyant L (2020) Strong differentiation within Diplocarpon rosae strains based on microsatellite markers and greenhouse-based inoculation protocol on Rosa. Plant Pathol 69:1093–1107. 10.1111/ppa.13182

McGregor CE, Miano DW, LaBonte DR, Hoy M, Clark CA, Rosa GJM (2009) Differential Gene Expression of Resistant and Susceptible Sweetpotato Plants after Infection with the Causal Agents of Sweet Potato Virus Disease. J Amer Soc Hort Sci 134:658–666. 10.21273/JASHS.134.6.658

Moore EM, Lopez Arias DC, Zlesak DC, Clark M, Yong D, Sproul J, Hokanson SC (2023) Identification of a Novel Rose Black Spot Disease Resistance Locus (Rdr6) in a Tetraploid Baby Love^TM^ Derived Population. horts 58:S1–S387. 10.21273/HORTSCI.58.9S.S1

Neu E, Domes HS, Menz I, Kaufmann H, Linde M, Debener T (2019) Interaction of roses with a biotrophic and a hemibiotrophic leaf pathogen leads to differences in defense transcriptome activation. Plant Mol Biol 99:299–316. 10.1007/s11103-018-00818-2

Neu E, Featherston J, Rees J, Debener T (2017) A draft genome sequence of the rose black spot fungus Diplocarpon rosae reveals a high degree of genome duplication. PLOS ONE 12:e0185310. 10.1371/journal.pone.0185310

Ngou BPM, Ding P, Jones JDG (2022) Thirty years of resistance: Zig-zag through the plant immune system. The Plant Cell 34:1447–1478. 10.1093/plcell/koac041

Patro R, Duggal G, Love MI, Irizarry RA, Kingsford C (2017) Salmon provides fast and bias-aware quantification of transcript expression. Nat Methods 14:417–419. 10.1038/nmeth.4197

Poland JA, Balint-Kurti PJ, Wisser RJ, Pratt RC, Nelson RJ (2009) Shades of gray: the world of quantitative disease resistance. Trends in Plant Science 14:21–29. 10.1016/j.tplants.2008.10.006

R Core Team (2024) R: A language and environment for statistical computing

Rawandoozi ZJ, Young EL, Yan M, Noyan S, Fu Q, Hochhaus T, Rawandoozi MY, Klein PE, Byrne DH, Riera-Lizarazu O (2022) QTL mapping and characterization of black spot disease resistance using two multi-parental diploid rose populations. Horticulture Research 9:uhac183. 10.1093/hr/uhac183

Rech SS, Heidt S, Requena N (2013) A tandem Kunitz protease inhibitor (KPI106)-serine carboxypeptidase (SCP1) controls mycorrhiza establishment and arbuscule development in Medicago truncatula. Plant J 75:711–725. 10.1111/tpj.12242

Ren J, Chen L, Liu J, Zhou B, Sha Y, Hu G, Peng J (2024) Transcriptomic insights into the molecular mechanism for response of wild emmer wheat to stripe rust fungus. Front Plant Sci 14. 10.3389/fpls.2023.1320976

Rouet C, Lee EA, Banks T, O’Neill J, LeBlanc R, Somers DJ (2020) Identification of a polymorphism within the Rosa multiflora muRdr1A gene linked to resistance to multiple races of Diplocarpon rosae W. in tetraploid garden roses (Rosa × hybrida). Theoretical and Applied Genetics 133:103–117. 10.1007/s00122-019-03443-9

Silva NF, Goring DR (2002) The proline-rich, extensin-like receptor kinase-1 (PERK1) gene is rapidly induced by wounding. Plant Mol Biol 50:667–685. 10.1023/a:1019951120788

Song J, Chen F, Lv B, Guo C, Yang J, Guo J, Huang L, Ning G, Yang Y, Xiang F (2024) Comparative transcriptome and metabolome analysis revealed diversity in the response of resistant and susceptible rose (Rosa hybrida) varieties to Marssonina rosae. Front Plant Sci 15. doi: 10.3389/fpls.2024.1362287

Soufflet-Freslon V, Marolleau B, Thouroude T, Chastellier A, Pierre S, Bellanger MN, Le Cam B, Bonneau C, Porcher L, Leclere A, Robert F, Felix F, Foucher F, Hibrand-Saint Oyant L (2019) Development of tools to study rose resistance to black spot. Acta Horticulturae 1232:213–220. 10.17660/ActaHortic.2019.1232.31

St Clair DA (2010) Quantitative Disease Resistance and Quantitative Resistance Loci in Breeding. Annual Review of Phytopathology 48:247–268. 10.1146/annurev-phyto-080508-081904

Su K, Zhao W, Lin H, Jiang C, Zhao Y, Guo Y (2023) Candidate gene discovery of Botrytis cinerea resistance in grapevine based on QTL mapping and RNA-seq. Front Plant Sci 14. 10.3389/fpls.2023.1127206

The UniProt Consortium (2023) UniProt: the Universal Protein Knowledgebase in 2023. Nucleic Acids Research 51:D523–D531. 10.1093/nar/gkac1052

Tian T, Liu Y, Yan H, You Q, Yi X, Du Z, Xu W, Su Z (2017) agriGO v2.0: a GO analysis toolkit for the agricultural community, 2017 update. Nucleic Acids Research 45:W122–W129. 10.1093/nar/gkx382

Voigt CA, Somerville SC (2009) Callose in Biotic Stress (Pathogenesis). In: Bacic A, Fincher F, Stone B (eds) Chemistry, Biochemistry, and Biology of 1-3 Beta Glucans and Related Polysaccharides. Elsevier, pp 525–562

von Malek B, Debener T (1998) Genetic analysis of resistance to blackspot (Diplocarpon rosae) in tetraploid roses: Theor Appl Genet 96:228–231. 10.1007/s001220050731

Waliczek TM, Byrne DH, Holeman DJ (2015) Growers’ and consumers’ knowledge, attitudes and opinions regarding roses available for purchase. Acta Horticulturae 235–239. 10.17660/ActaHortic.2015.1064.26

Wang G, Ellendorff U, Kemp B, Mansfield JW, Forsyth A, Mitchell K, Bastas K, Liu C-M, Woods-Tör A, Zipfel C, de Wit PJGM, Jones JDG, Tör M, Thomma BPHJ (2008) A Genome-Wide Functional Investigation into the Roles of Receptor-Like Proteins in Arabidopsis. Plant Physiology 147:503–517. 10.1104/pp.108.119487

Wang W, Devoto A, Turner JG, Xiao S (2007) Expression of the Membrane-Associated Resistance Protein RPW8 Enhances Basal Defense Against Biotrophic Pathogens. MPMI 20:966–976. 10.1094/MPMI-20-8-0966

Whitaker VM, Bradeen JM, Debener T, Biber A, Hokanson SC (2010a) Rdr3, a novel locus conferring black spot disease resistance in tetraploid rose: genetic analysis, LRR profiling, and SCAR marker development. Theor Appl Genet 120:573–585. 10.1007/s00122-009-1177-0

Whitaker VM, Debener T, Roberts AV, Hokanson SC (2010b) A standard set of host differentials and unified nomenclature for an international collection of Diplocarpon rosae races: Diplocarpon rosae races. Plant Pathology 59:745–752. 10.1111/j.1365-3059.2010.02281.x

Wickham H (2016) ggplot2. Springer International Publishing, Cham

Wu Z, Li M, Dong OX, Xia S, Liang W, Bao Y, Wasteneys G, Li X (2019) Differential regulation of TNL-mediated immune signaling by redundant helper CNLs. New Phytologist 222:938–953. 10.1111/nph.15665

Xiao S, Jiang L, Wang C, Ow DW (2021) Arabidopsis OXS3 family proteins repress ABA signaling through interactions with AFP1 in the regulation of ABI4 expression. Journal of Experimental Botany 72:5721–5734. 10.1093/jxb/erab237

Yang H, Li Y, Hua J (2006) The C2 domain protein BAP1 negatively regulates defense responses in Arabidopsis. The Plant Journal 48:238–248. 10.1111/j.1365-313X.2006.02869.x

Yang H, Yang S, Li Y, Hua J (2007) The Arabidopsis BAP1 and BAP2 Genes Are General Inhibitors of Programmed Cell Death. Plant Physiology 145:135–146. 10.1104/pp.107.100800

Yang S, Xu T, Yang Y, Pei W, Luo L, Yu C, Wang J, Cheng T, Zhang Q, Pan H (2022) H2O2 accumulation plays critical role in black spot disease resistance in roses. Hortic Environ Biotechnol. 10.1007/s13580-022-00458-y

Yang Y, Zhang Y, Ding P, Johnson K, Li X, Zhang Y (2012) The Ankyrin-Repeat Transmembrane Protein BDA1 Functions Downstream of the Receptor-Like Protein SNC2 to Regulate Plant Immunity. Plant Physiology 159:1857–1865. 10.1104/pp.112.197152

Yu M-H, Zhao Z-Z, He J-X (2018) Brassinosteroid Signaling in Plant–Microbe Interactions. International Journal of Molecular Sciences 19:4091. 10.3390/ijms19124091

Zeilmaker T, Ludwig NR, Elberse J, Seidl MF, Berke L, Van Doorn A, Schuurink RC, Snel B, Van den Ackerveken G (2015) DOWNY MILDEW RESISTANT 6 and DMR6-LIKE OXYGENASE 1 are partially redundant but distinct suppressors of immunity in Arabidopsis. Plant J 81:210–222. 10.1111/tpj.12719

Zheng X, Long Y, Liu X, Han G, Geng X, Ju X, Chen W, Xu T, Tang N (2024) RcWRKY40 regulates the antagonistic SA–JA pathway in response to Marssonina rosae infection. Scientia Horticulturae 332:113178. 10.1016/j.scienta.2024.113178

Zlesak DC (2007) Rose Rosa x hybrida. In: Anderson NO (ed) Flower Breeding and Genetics. pp 695–740

Zlesak DC, Ballantyne D, Holen M, Clark A, Hokanson SC, Smith K, Zurn JD, Bassil NV, Bradeen JM (2020) An Updated Host Differential Due to Two Novel Races of Diplocarpon rosae Wolf, the Causal Agent of Rose Black Spot Disease. HortScience 55:1756–1758. 10.21273/HORTSCI14902-20

Zurn JD, Zlesak DC, Holen M, Bradeen JM, Hokanson SC, Bassil NV (2018) Mapping a Novel Black Spot Resistance Locus in the Climbing Rose Brite Eyes^TM^ (‘RADbrite’). Frontiers in Plant Science 9. 10.3389/fpls.2018.01730

Zurn JD, Zlesak DC, Holen M, Bradeen JM, Hokanson SC, Bassil NV (2020) Mapping the black spot resistance locus Rdr3 in the shrub rose ‘George Vancouver’ allows for the development of improved diagnostic markers for DNA-informed breeding. Theoretical and Applied Genetics 133:2011–2020. 10.1007/s00122-020-03574-4

